# Functional and structural asymmetry suggest a unifying principle for catalysis in integral membrane-bound pyrophosphatases

**DOI:** 10.1101/2022.11.07.515396

**Authors:** Jannik Strauss, Craig Wilkinson, Keni Vidilaseris, Orquidea Ribeiro, Jianing Liu, James Hillier, Anssi Malinen, Bernadette Gehl, Lars J.C. Jeuken, Arwen R. Pearson, Adrian Goldman

**Author notes:** Numaferm GmbH, Düsseldorf, Germany (Jannik Strauss, James Hillier). Department of Applied Physics, Aalto University, FI-00076, AALTO, Finland.

## Abstract

Membrane-bound pyrophosphatases (M-PPases) are homodimeric primary ion pumps that couple the transport of Na^+^- and/or H^+^ across membranes to the hydrolysis of pyrophosphate. Their role in the virulence of protist pathogens like *Plasmodium falciparum* makes them an intriguing target for structural and functional studies. Here, we show the first structure of a K^+^-independent M-PPase, asymmetric and time-dependent substrate binding in time-resolved structures of a K^+^-dependent M-PPase, and demonstrate pumping-before-hydrolysis by electrometric studies. We suggest how key residues in helix 12, 13, and the exit channel loops affect ion selectivity and K^+^-activation due to a complex interplay of residues that are involved in subunit-subunit communication. Our findings not only explain ion selectivity in M-PPases but also why they display half-of-the-sites reactivity. Based on this we propose, for the first time, a unified and testable model for ion pumping, hydrolysis, and energy-coupling in *all* M-PPases, including those that pump both Na^+^ and H^+^.

## Introduction

Pyrophosphatases (PPases) catalyse the hydrolysis of inorganic pyrophosphate (PP_i_), a by- product of nearly 200 biosynthetic reactions across all kingdoms of life (Heinonen, 2001; Lahti, 1983). Soluble PPases (S-PPases) are responsible for recycling the intracellular PP_i_ pool in all types of organisms, whereas the function of membrane-bound PPases (M-PPases) extends beyond mere PP_i_ hydrolysis (H. Baltscheffsky et al., 1966; Moyle et al., 1972). They utilise the energy stored in the phosphoanhydride bond of PP_i_ by coupling its hydrolysis to the directed transport of sodium ions (Na^+^) and/or protons (H^+^) across membranes, but are only present in plants, parasitic protists, and certain prokaryotes (M. Baltscheffsky et al., 1999; Luoto, Baykov, et al., 2013; Malinen et al., 2007). They are classified into different subclasses based on their ion pumping selectivity and co-factor dependence. To date, H^+^-pumping (H^+^-PPase), Na^+^- pumping (Na^+^-PPase) and dual-pumping (Na^+^,H^+^-PPase) M-PPases have been found (Luoto, Baykov, et al., 2013; Nordbo et al., 2016), most of which require (K^+^) for maximal catalytic activity. Only in H^+^-PPases has evolution given rise to a subclass of K^+^-independent enzymes (Table 1)(H. Baltscheffsky et al., 1966; Walker & Leigh, 1981).

**Table 1:** M-PPase classification into different subclasses.

| Cation pumping specificity | Monovalent cation dependence | Residue at |  | Semi-conserved glutamate | Structural data (PDB) | Example | Reference |
| --- | --- | --- | --- | --- | --- | --- | --- |
|  |  | 12.46 | 12.49 |  |  |  |  |
| Na<sup>+</sup> | K<sup>+</sup>, Na<sup>+</sup> | A | G | 6.53 | 4AV3, 4AV6, 5LZQ, 5LZR, 6QXA | *Thermotoga maritima* | (Kellosalo et al., 2012; Li et al., 2016; Vidilaseris et al., 2019) |
| H<sup>+</sup> | K<sup>+</sup> | A | T | 6.57 | 4A01, 5GPJ, 6AFS, 6AFT, 6AFU, 6AFV, 6AFW, 6AFX, 6AFY, 6AFZ | *Vigna radiata* | (Li et al., 2016; Lin et al., 2012; Tsai et al., 2019) |
|  | - | K | G | 6.53 | This study | *Pyrobaculum aerophilum* | - |
| Na<sup>+</sup>,H<sup>+</sup> (dual) | K<sup>+</sup>, Na<sup>+</sup> | A | G | 6.53 | - | *Clostridium leptum* | - |

### M-PPases in human health and global food security

Plants, certain prokaryotes and parasitic protists utilise the PP_i_ pool as an additional energy source to survive low-energy and high-stress conditions by establishing electrochemical gradients across membranes (García-Contreras et al., 2004; Lander et al., 2016). This makes M-PPases, which could be of benefit in the fight against existing and emerging challenges to global food security and human health, a valuable target for structural and functional studies. For example, overexpression of M-PPases improves drought tolerance in various transgenic plants (Esmaeili et al., 2019; Gaxiola et al., 2001; Park et al., 2005); this could be of great importance, as global warming induces crop losses of 3.2-7.4 % per degree rise in the global mean temperature (Lesk et al., 2016; Zhao et al., 2017; Ray et al., 2019). Global warming also facilitates the pole-wards spread of insect vectors for parasites such as *Plasmodium falciparum*, putting millions of new people at risk of life-threatening diseases in the coming decades (Hertig, 2019; Ryan et al., 2019). Consequently, impairing cellular homoestasis in protozoan parasites that harbor M-PPases, such as *Plasmodium* ssp. (malaria), *Leishmania spp*. (leishmaniasis), *Trypanosome spp.* (trypanosomiasis) and *Toxoplasma gondii* (toxoplasmosis) is a promising approach to combating these dieases (Zhang et al., 2018; Lemercier et al., 2002; Liu et al., 2014). Recently, we have developed non-phosphorus M-PPase inhibitors with the potential for further development into therapeutic molecules (Johansson et al., 2020, 2021; Vidilaseris et al., 2019).

### Structural features of M-PPases

M-PPases are large (66-89 kDa), single-domain integral membrane proteins comprised of two identical monomers, each with 15-17 transmembrane helices (Luoto et al., 2015). To date, the structures of only two are known, a K^+^-dependent H^+^-PPase from *Vigna radiata* (*Vr*PPase) and a K^+^-dependent Na^+^-PPase from *Thermotoga maritima* (*Tm*PPase), with structures available for both in various catalytic states (Kellosalo et al., 2012; Li et al., 2016; Lin et al., 2012; Tsai et al., 2019; Vidilaseris et al., 2019). In general, the helices of each subunit arrange into an inner ring (helices 5, 6, 11, 12, 15, 16) containing the functional core (active site, coupling funnel, ion gate, exit channel) and an outer ring (helices 1-4, 7-10, 13, 14) of largely unknown function (Figure 1A). In the following, we use the residue numbering scheme X^Y.Z^ in which X represents the amino acid as single-letter code, Y denotes the helix on which it is located and Z defines the offset of a well conserved in the centre of this helix (Ballesteros & Weinstein, 1995). This simplifies residue comparison between proteins and highlights conservation. A translation to conventional residue numbering can be found in Supplementary table 1.

**Figure 1:**
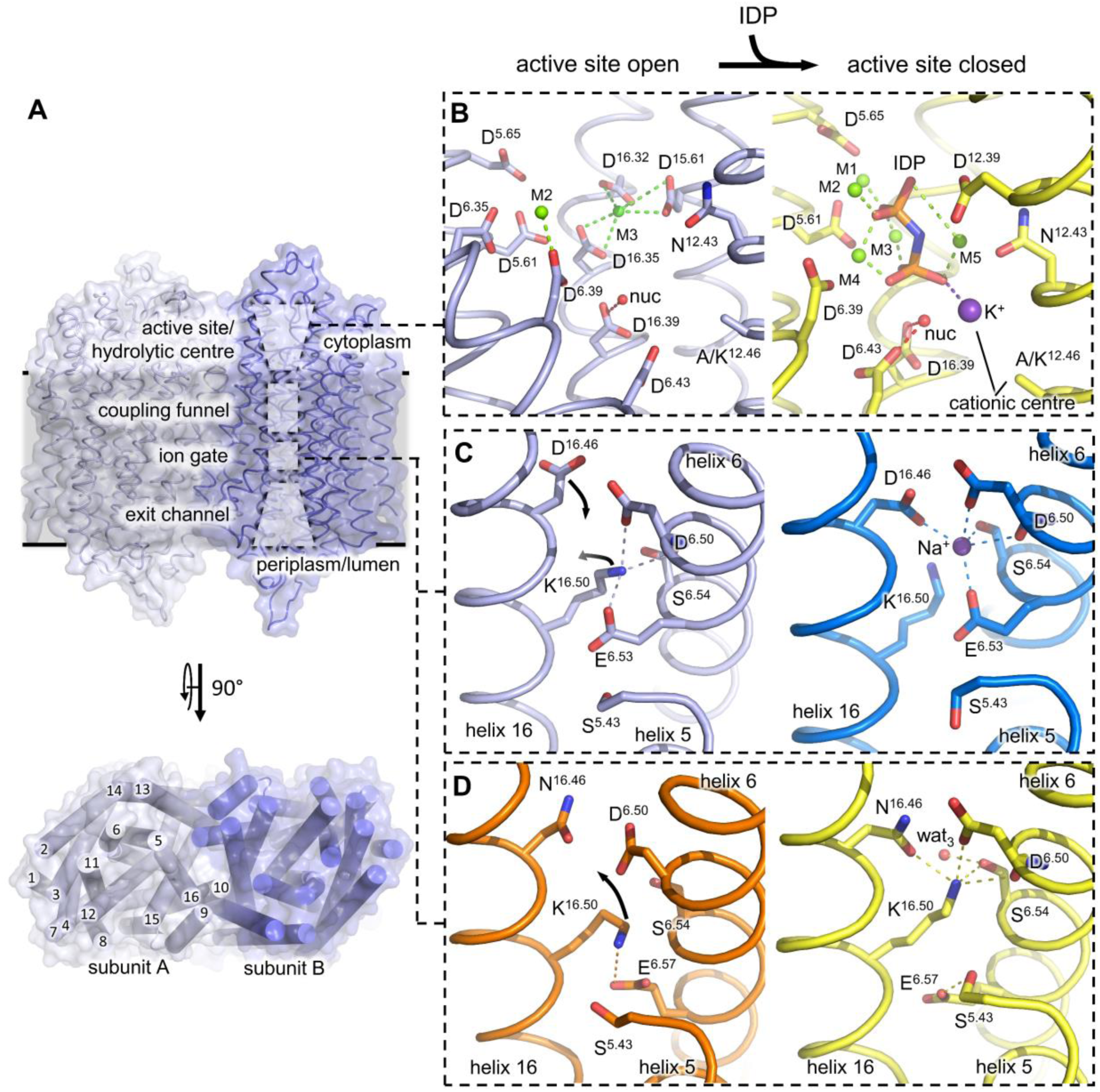
Structural features of M-PPases. Protein colouring follows previous publications (Li et al., 2016; Vidilaseris et al., 2019) that used shades of yellow/ orange for *Vr*PPase and shades of blue for *Tm*PPase structures. (**A**) Homodimeric M-PPase viewed from the membrane plane (top) with the functional core (active site, coupling funnel, ion gate, exit channel) highlighted by dashed boxes. Concentric ring arrangements of transmembrane helices (bottom) viewed from the cytoplasmic site. Loops were removed for clarity. (**B**) Close-up view of the *Tm*PPase active site (resting-state: left panel, active-state: right panel) with helix 11 removed for clarity. M1-5 is Mg^2+^ (active-state), M3 is inhibitory Ca^2+^ (resting-state). K^+^ (purple sphere) is part of the cationic centre in K^+^- dependent M-PPases (with A^12.46^). The non-hydrolysable substrate analogue IDP (imidodiphosphate) is shown in orange. (**C-D**) Close-up view of the ion gate in K^+^-dependent Na^+^-PPases (**C**) or K^+^-independent H^+^-PPases (**D**). Left panels show the residue orientation when the active site is open, right panel shows the residue configuration when the active site is fully closed upon IDP-binding to the active site. Na^+^ shown as blue sphere, structural water (wat3) displayed as red sphere. Dashed lines highlight key interactions.

#### Active site

The active site in M-PPases protrudes about 20 Å above the membrane plane into the cytoplasm (Figure 1A), containing completely conserved Asp, Asn and Lys residues that provide the basis for PP_i_ binding and hydrolysis (Kellosalo et al., 2012; Lin et al., 2012). Amino acid substitutions in this region typically lead to inactive protein (Lin et al., 2012; Nakanishi et al., 2001; Schultz & Baltscheffsky, 2003; Hirono et al., 2007; Asaoka et al., 2014). The aspartate and asparagine side chains coordinate up to five Mg^2+^, which capture PP_i_ in a metal cage, whereas lysine side chains directly stabilise PP_i_ binding at the active site (Figure 1B). Of the up to five Mg^2+^ present at the active site, two are brought in by the enzymatically active substrate (Mg_2_PP_i_), two bind to activating high-affinity sites (*K*_d_: ∼20-460 µM) and one binds to a putatively inhibitory low-affinity site (*K*_d_: ∼100 mM)(Maeshima, 2000; Malinen et al., 2008). A nucleophilic water molecule is poised to attack PP_i_ for reaction initiation and interacts with either one (D^6.43^, resting state) or two aspartates (D^6.43^ and D^16.39^, active state)(Figure 1B).

#### Coupling funnel

The coupling funnel links the active site to the ion gate in the centre of the membrane (Figure 1A) and couples PP_i_ hydrolysis to the transport of Na^+^, H^+^ or both across the membrane (Kellosalo et al., 2012; Lin et al., 2012). A set of highly conserved charged residues including R/Q^5.50^, D^6.43^, D^6.50^, D/S^11.50^, K^12.50^, K^16.38^ and D^16.39^ are arranged to form a Grotthus chain through the top half of the membrane, allowing ion translocation (Kellosalo et al., 2012; Lin et al., 2012). Of these residues, D^6.43^ and D^16.39^ sit at the interface of the active site (Figure 1B), whereas D^6.50^ connects to the ion gate (Figure 1C).

#### Ion gate and exit channel

The ion gate functions as an ion selectivity filter for pumping in the membrane spanning protein region. A set of four residues, E^6.53/57^, D^6.50^, S^6.54^ D/N^16.46^, form a Na^+^/H^+^ binding site. Binding of Mg_2_PP_i_ to the active site requires a downward shift of helix 12 and corkscrew motion at helix 6 and 16, which affects the ion gate configuration of K^+^-dependent Na^+^-PPases (*Tm*PPase) and K^+^-dependent H^+^-PPases (*Vr*PPase) differently. In K^+^-dependent Na^+^-PPases, K^16.50^ rotates out of the Na^+^-binding site, making it available for ion binding (Figure 1C). In K^+^-dependent H^+^-PPases, K^16.50^ reorientation unmasks a proton binding site instead. The K^16.50^-E^6.57^ ion-pair breaks and a D/N^16.46^- K^16.50^-D^6.50^-S^6.54^ interaction forms, which leaves the side chain of E^6.57^ stabilised only by a hydrogen bond to S^5.43^ in a hydrophobic protein environment (Figure 1D). It has been proposed that E^6.57^ is protonated, thus linking structural differences at the ion gate between K^+^-dependent Na^+^-PPases and K^+^-dependent H^+^- PPases to the observed ion pumping selectivity (Li et al., 2016). The exit channel below the ion gate has low sequence conservation but its properties are important in facilitating ion release (Tsai et al., 2019).

#### Dimer interface

The interface between monomers is formed by residues of the outer ring helices 10, 13 and inner ring helix 15 (Figure 1A) that interact with the opposing subunit *via* hydrogen bonds and hydrophobic interactions (Kellosalo et al., 2012; Lin et al., 2012). The dimer interface has not typically been considered as key to the function of M-PPases as all of the catalytic machinery seems to be located in a single subunit (Kellosalo et al., 2012; Lin et al., 2012), but a growing body of structural and functional evidence points to (1) that M-PPases are functionally asymmetric, and (2) that the dimer interface mediates key inter-subunit interactions (Anashkin et al., 2021; Artukka et al., 2018; Vidilaseris et al., 2019) through coupled helix motions during the catalytic cycle(Li et al., 2016).

### Energy coupling

The chronological order of PP_i_ hydrolysis and ion pumping is a point of active discussion for M-PPases (Baykov, 2020; Li et al., 2016). The two opposing mechanisms of energy coupling either postulate “pumping-before-hydrolysis” or “pumping-after-hydrolysis”. The “pumping- after-hydrolysis” model, also called “Mitchell-direct”, postulates that PP_i_ hydrolysis and ion pumping occur simultaneously and that the H^+^ released from the nucleophilic water during PP_i_ hydrolysis is the one pumped after *n* cycles, where *n* is the number of downstream ion binding sites (Baykov, 2020). This was extended by a “billiard-type” mechanism to explain Na^+^-transport in which the generated H^+^ pushes Na^+^ into the exit channel for pumping (Baykov et al., 2013). In contrast, the “pumping-before-hydrolysis” model, also called “binding-change”, favours a mechanism in which ion pumping precedes hydrolysis and is initiated by the closure of the active site and associated helical rearrangements, explaining both H^+^ and Na^+^ pumping (Li et al., 2016). The transported ion may originate from the medium or preceding hydrolysis events and can explain both H^+^- and Na^+^-pumping. The overall negative charge at the ion gate that results from “pumping-before-hydrolysis” would then promote the abstraction of a H^+^ from nucleophilic water at the active site and thereby drive the hydrolysis of PP_i_. The generated H^+^ could enter the Grotthus chain and reset the ion gate (Li et al., 2016).

### The evolution of K^+^-independence and ion pumping selectivity

Two coupled changes are correlated with the evolution of K^+^-independent H^+^-PPases: A^12.46^K and G/A^12.49^T (Belogurov & Lahti, 2002). Of them, the A/K^12.46^ change is the one that defines K^+^-dependence (Artukka et al., 2018): the ε-NH_3_^+^ of K^12.46^ has been postulated to replace K^+^ in the cationic centre both functionally and structurally (Figure 1A). However, there has been no structural data available to support this idea. Moreover, although the A^12.46^K and G/A^12.49^T changes are tightly coupled evolutionarily, there is no functional role so far ascribed to the residue at position 12.49 (Belogurov & Lahti, 2002). It appears to be involved in K^+^-binding as A^12.49^T single variants of K^+^-dependent H^+^-PPases show a three-fold reduced affinity for K^+^, but it remains unclear how changes at this position affect the cationic centre, which is ∼10 Å away (Belogurov & Lahti, 2002). Alternatively, G/A/T^12.49^ may play a crucial role in substrate inhibition as this regulatory mechanism is lost in M-PPases when interfering with the authentic state of the cationic centre, *e.g.* in A^12.49^K single variants of K^+^-dependent H^+^-PPases (Artukka et al., 2018).

In contrast to K^+^-dependence, there are no conserved residue patterns that correlate with ion pumping selectivity across *all* M-PPase subclasses, but the C-terminal shift of a key glutamate at the ion gate of *K^+^-dependent* M-PPases by one helix turn (E^6.53è57^) is coupled to a change in selectivity (Na^+^èH^+^)(Lin et al., 2012). The transition from Na^+^ to H^+^ pumping in this model simply requires the repositioning of a single residue without the need of a mechanistic change (Li et al., 2016). When the semi-conserved glutamate is located one helix turn down, K^16.50^ continues to block the Na^+^-binding site upon substrate binding at the active site, while E^6.57^ reorientates and can now accommodate a proton (Figure 1C). However, this model fails to explain ion pumping selectivity in K^+^-independent H^+^-PPases or K^+^-dependent Na^+^,H^+^-PPases, as both contain E^6.53^ (Table 1). It might make more sense to consider E^6.53^ the conserved position, with mutations in K^+^-dependent H^+^-PPases containing E^6.57^.

There are thus clearly unanswered questions: what is the mechanism of energy coupling and of ion pumping selectivity; what is the structure of K^+^-independent M-PPases; and what is the structural/functional basis of catalytic asymmetry. To address these questions, we solved the first structure of a K^+^-independent M-PPase (from the thermophile *Pyrobaculum aerophilum*, *Pa*PPase), performed enzymatic assays on native *Pa*PPase and three variants (A^12.46^K and A^12.49^T, and the double mutant), and conducted electrometric as well as time-resolved crystallographic studies on *Tm*PPase, a K^+-^dependent Na^+^-PPase. These data provide a structural mechanism for half-of-the-sites reactivity in M-PPases. By *requiring* a dimer for a complete catalytic cycle, our new model for M-PPase catalysis suggests a resolution to the “binding-change” *versus* “Mitchell-direct” controversy in energy coupling.

## Results

### Structure of *Pa*PPase

Purified wild-type *Pa*PPase (Figure 2-figure supplement 1A-B) readily crystallised in vapour-diffusion set-ups, but despite extensive optimisation efforts the diffraction was anisotropic. The data were submitted to the STARANISO webserver and the structure was solved by molecular replacement (MR) using a modified *Tm*PPase:Mg_5_IDP (PDB: 5LZQ) search model with all loops and hetero atoms removed. This yielded a structure with one *Pa*PPase homodimer molecule per asymmetric unit and resolutions of 5.3 Å, 4.1 Å and 3.8 Å along h, k and l, respectively (Supplementary table 2). After initial refinement, positive mF_o_-dF_c_ density was observed at 3 σ in both subunits of the *Pa*PPase active site, at the ion gate (Figure 2-figure supplement 2A) and in the dimer interface. We built a Mg_5_IDP complex into the active site, as seen in other IDP-bound M-PPase structures, and two water molecules in regions with excess positive mF_o_-dF_c_ density that bridge between loop_5-6_ and the metal cage (Figure 2-figure supplement 2B). A structural water was built into the positive mF_o_-dF_c_ density at the ion gate as in the high resolution *Vr*PPase structure and a sulfate molecule (SO_4_^2-^) was placed at the dimer interface (Figure 2-figure supplement 2B). The electron density maps improved throughout refinement and the final R_work_/R_free_ was 28.9%/31.1% with appropriate stereochemistry for this resolution range (3.8-5.3 Å).

**Figure 2:**
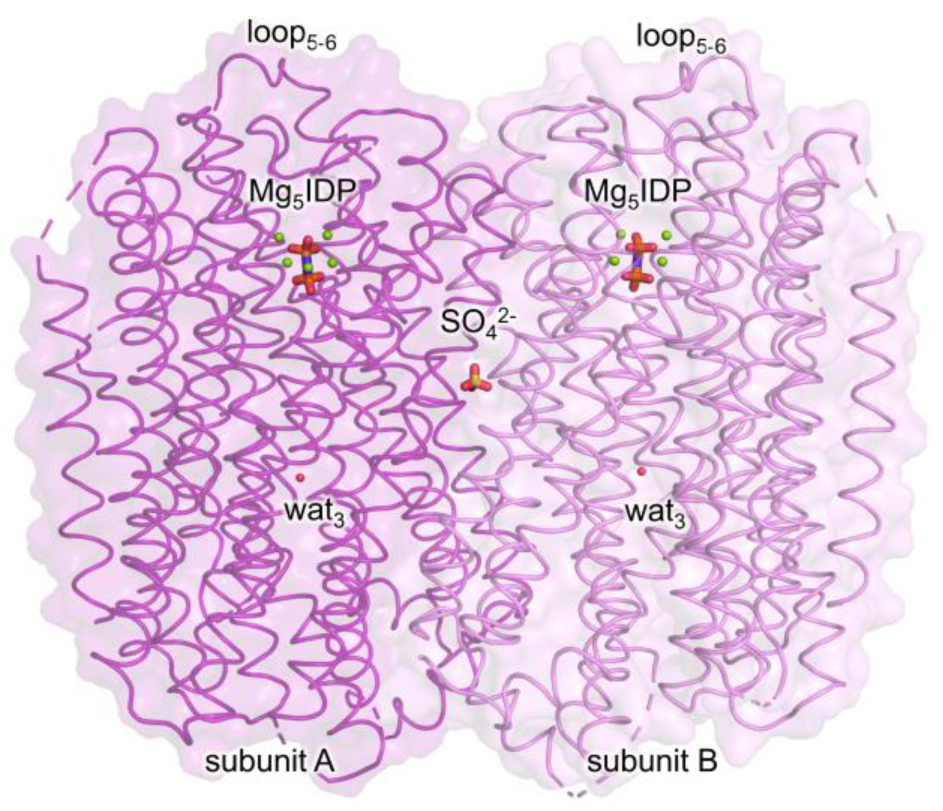
Overview of the *Pa*PPase:Mg5IDP structure. Subunits, loop5-6 and ligands/structural water molecules are annotated.

### Structural overview and comparison of M-PPase structures

The *Pa*PPase structure is in the Mg_5_IDP-bound state with loop_5-6_ closed and a structural water located at the ion gate (Figure 2). In what follows, structural alignments and root mean square deviation (r.m.s.d.) calculations are based on the Cα atom of subunit A (both subunits are nearly identical; r.m.s.d./C_α_: 0.27 Å), unless stated otherwise. The overall structure is very similar to other published M-PPase structures with an average r.m.s.d 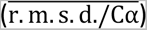 of 1.37 ±0.18 Å to IDP-bound structures, 1.41 ±0.15 Å to product-bound structures, and a r.m.s.d./C_α_ of 1.67 Å to the resting state structure (Supplementary table 3). In general, outer ring helices display more variability 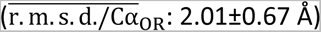 than inner-ring helices 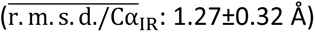 when compared to *Tm*/*Vr*PPase:Mg_5_IDP.

Alignment-independent inter atom difference distance matrices (DiDiMa) highlighted outer ring helices 13-14 (and 2-3 in *Vr*PPase but not *Tm*PPase) as regions with major structural differences when comparing identical enzyme states (Figure 3-figure supplement 1A). In previously solved IDP-bound structures, helices 13-14 are consistently bent halfway through the membrane by about 9° to remain near the cytoplasmic regions of helix 5 (Figure 3-figure supplement 1B). This enables propagation of motions from the inner to the outer ring and into the other subunit (indicated by apostrophe) *via* E^5.71^–R^13.62^–R/I/K^10.33ˈ^(position at 10.33 not conserved in Vr/*Tm*/*Pa*PPase) (Figure 3-figure supplement 2A-B) and was linked to loop_5-6_ and subsequent helical rearrangements (Li et al., 2016). In our new structure, the E^5.71^-R^13.62^-R/I/K^10.33ˈ^ interaction is lost (Figure 3-figure supplement 1C): the cytoplasmic regions of helices 13-14 are straightened (Figure 3-figure supplement 1B), resembling resting-state (Figure 3-figure supplement 1C) and product-bound (Figure 3-figure supplement 1D-E) structures despite having IDP bound. This suggests a different role for helix 13-14 movement in M-PPase function than previously thought (see Discussion).

**Figure 3:**
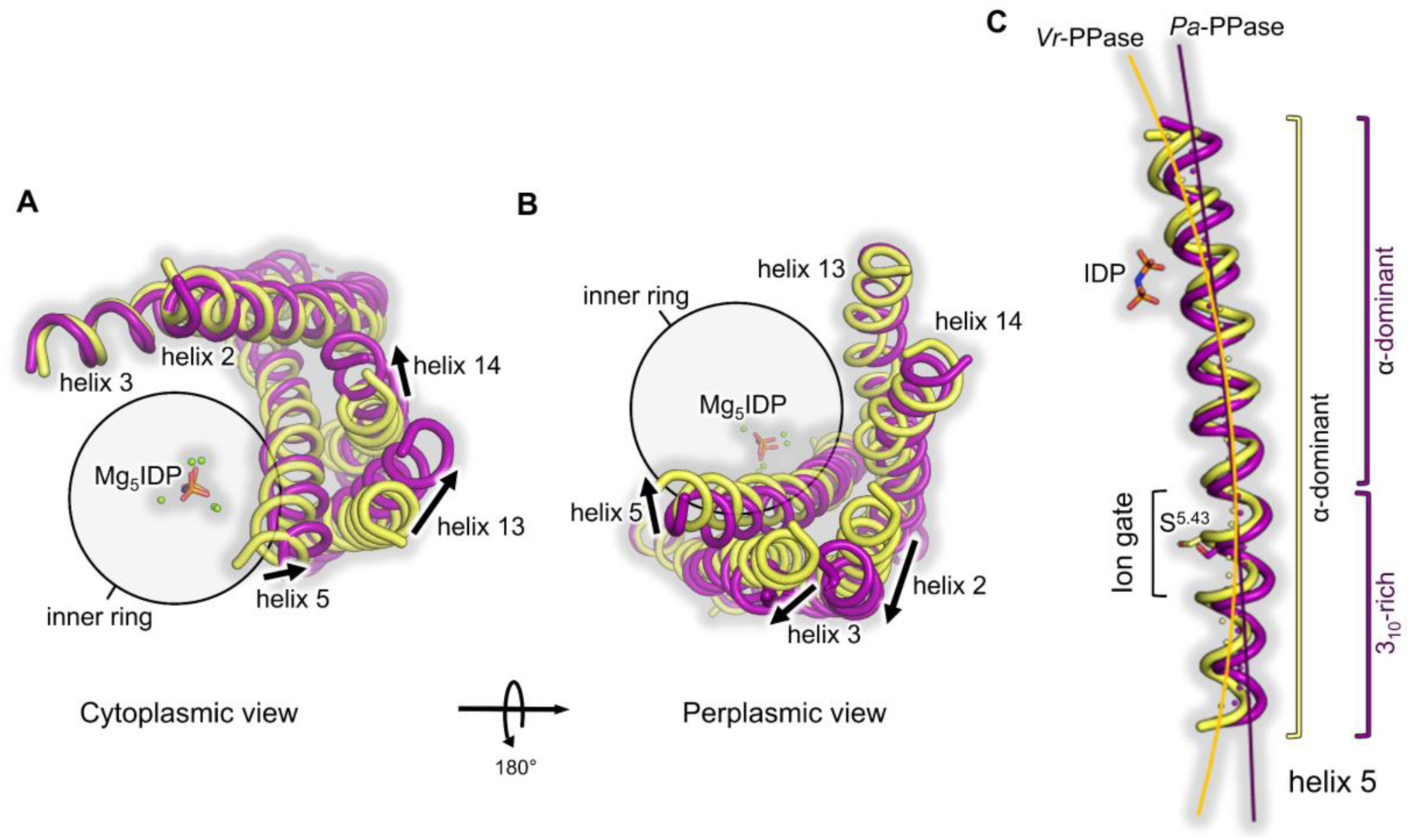
Comparison of helices 2-3, 5, and 13-14 orientations in *Pa*PPase:Mg5IDP to *Vr*PPase:Mg5IDP. *Pa*PPase:Mg5IDP is shown in purple and *Vr*PPase:Mg5IDP is shown in yellow. Major conformational changes are indicated by arrows. (**A**) Close-up view of helix 5 showing its straightening in *Pa*PPase:Mg5IDP compared to *Vr*PPase:Mg5IDP. Helix straightening is highlighted by a curve running through the centre of helix 5, which was manually fitted to the local helix origin points that were obtained from HELANAL-Plus analysis (displayed as spheres in the helix centre). (**B**) Straigtheing of helix 5 pushes helices 13-14 away from the inner ring on the cytoplasmic site. (**C**) Straigtheing of helix 5 pushes of helix 2-3 away from the inner ring on the periplasmic site.

The only inner ring (IR) helix with above-average conformational differences to previously published IDP-bound structures (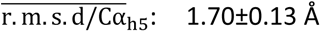 *versus* 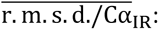 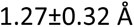) is helix 5, around which outer ring helices 2-3 and 13-14 cluster. Helix 5 is straighter than in other M-PPases (Figure 3A, Supplementary table 4), which also straightens helices 13-14 (cytoplasmic side, Figure 3B) and 2-3 (periplasmic side, Figure 3C) by pushing them away from the inner ring. Additionally, helix 5 is more tightly wound in *Pa*PPase:Mg_5_IDP due to the presence of twice as many 3_10_ hydrogen bonds around S^5.43^ and towards its flanking periplasmic segment (Supplementary table 5). Consequently, the side chain orientations are different in this region compared with *Vr/Tm*PPase:Mg_5_IDP. This is particularly interesting, as S^5.43^ is part of the enzymatic core region defining ion selectivity, which, until now, could not be explained for K^+^-independent M-PPases.

### Structural and functional characterisation of K^+^ independence in *Pa*PPase

The coordination of the Mg_5_IDP complex (Figure 4A) at the active site by acidic residues is almost identical to *Tm*/*Vr*PPase:Mg_5_IDP (r.m.s.d./C_α_: 0.81/0.73 Å, alignment of catalytic residues in the active site of subunit A), and the hydrolytic pocket volume is about 1200 Å^3^ for all three structures. However, there are some interesting structural changes compared with previously solved K^+^-dependent M-PPases with A^12.46^ and A/G^12.49^:

**Figure 4:**
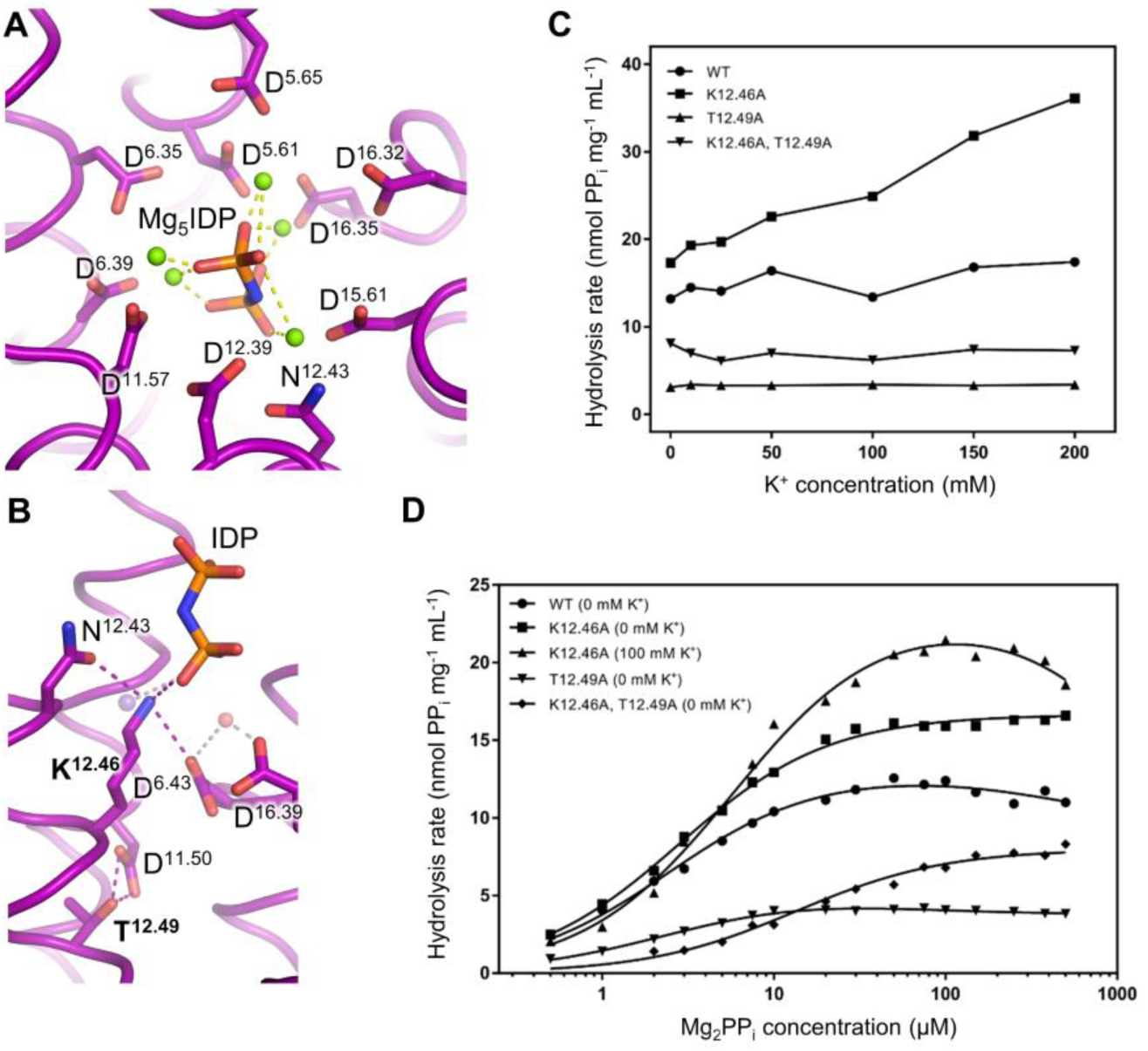
Structural overview and functional characterisation of the cationic centre in the *Pa*PPase active site. (**A**) Active site with IDP coordinated (dashed lines) in a Mg^2+^ metal cage (green spheres). (**B**) K^+^/K^12.46^ cationic centre with K^+^ (transparent purple sphere) and nucleophilic water (transparent red sphere) modelled into the structure based on its position in *Vr*PPase:Mg5IDP. Key residues of K^+^-independence are labelled in bold. Their interaction is shown by dashed lines. (**C**) Potassium dependency of PPi hydrolysis of wild-type and variant *Pa*PPase. (**D**) Wild-type and variant *Pa*PPase kinetics in the absence and presence of 100 mM K^+^. All data were collected in the presence of 5 mM free Mg^2+^. Wild-type (0 mM K^+^), K^12.46^A (100 mM K^+^), and T^12.49^A (0 mM K^+^) show the best fit to Equation 1, while K^12.46^A (0 mM K^+^) and K^12.46^A,T^12.49^A (0 mM K^+^) show the best fit to the Michaelis-Menten equation.

First, the side chain of K^12.46^ reaches into the hydrolytic pocket of *Pa*PPase. Residues with flexible side chains, such as the key players of K^+^-(in)dependence (K^12.46^) and ion selectivity (K^16.50^) had poor electron density in 2mF_o_-dF_c_ maps (Figure 2-figure supplement 3), so we modelled them by careful analysis of all possible rotamer conformations in the context of the local environment, taking into account hydrogen bonding and potential clashes (Supplementary table 6). The modelled conformation of K^12.46^ shows the smallest van der Waals (vdW) radii overlap to nearby atoms (0.62 Å), good hydrogen bonding (n_h-bonds_: 3) and avoids severe clashes. It coordinates IDP, replacing the K^+^ identified in *Vr*PPase:Mg_5_IDP (Figure **4**B) and hydrogen bonds N^12.43^ and D^6.43^. K^12.46^ thus substitutes for K^+^ in substrate coordination, explaining why *Pa*PPase does not require K^+^ for enzymatic activity. The two next best rotamers differ only in C_3_ and can also explain K^+^-independence as they remain near the K^+^-binding site and coordinate IDP. Second, T^12.49^ interacts with D^11.50^ in *Pa*PPase (Figure 4B), a direct result of the coupled A/G^12.49^T change in K^+^-independent M-PPases. This appears to lead to other coupled interactions with helix 6 that are nearby; in particular, the helical geometry around position 6.47 is changed (Supplementary table 5), also affecting the geometry at the catalytically essential general base D^6.50^.

We then measured the activity of wild-type, K^12.46^A, T^12.49^A and double variants. As expected, hydrolysis by wild-type enzyme is not activated by K^+^ (Figure 4C) but is inhibited by substrate (Figure 4D), with binding of the second PP_i_ very weak (Table 2). The K^12.46^A variant is weakly activated by K^+^ (Figure 4C). Without K^+^, the K^12.46^A variant displays Michaelis-Menten kinetics; in the presence of 100 mM K^+^, it has similar substrate inhibition as wild-type (Figure **4**D) except that V_2_ is now zero (Table 2). The T^12.49^A variant is essentially inactive (Figure 4C), while the double mutant no longer shows signs of substrate inhibition (Figure 4D): conventional Michaelis-Menten kinetics provide acceptable fits to the data (Table 2). Taken together, these suggest that helix 12, the site of the largest motion in the active site during catalysis with key residues K^12.46^ and T^12.49^ (Li et al., 2016), may play a crucial role in inter-subunit communication and, furthermore, that the observed half-of-the-sites reactivity (Vidilaseris et al., 2019; Artukka et al., 2018; Anashkin et al., 2021) may be key to understanding the true catalytic cycle (see Discussion).

**Table 2:** Kinetic parameters for PPi hydrolysis of PaPPase.

| Parameter | Wild-type<br>(0 mM $K^+$) | $K^{12.46}A$<br>(0 mM $K^+$) | Value<br>$K^{12.46}A$<br>(100 mM $K^+$) | $T^{12.49}A$<br>(0 mM $K^+$) | $K^{12.46}A, T^{12.49}A$<br>(0 mM $K^+$) |
| --- | --- | --- | --- | --- | --- |
| Equation | Substrate inhibition | Michaelis-Menten | Substrate inhibition | Substrate inhibition | Michaelis-Menten |
| $V_{max}$ (nmol PPI·mg<sup>-1</sup>·min<sup>-1</sup>) | | 16.64 ± 0.12 | | | 7.98 ± 0.21 |
| $V_1$ (nmol PPI·mg<sup>-1</sup>·min<sup>-1</sup>) | 12.89 ± 0.34 | | 23.34 ± 0.60 | 4.739 ± 0.12 | |
| $V_2$ (nmol PPI·mg<sup>-1</sup>·min<sup>-1</sup>) | 9.44 ± 3.22 | | 0 | 3.78 ± 0.094 | |
| $K_m$ (μM) | | 2.81 ± 0.08 | | | 13.48 ± 1.12 |
| $K_{m1}$ (μM) | 2.42 ± 0.21 | | 5.86 ± 0.39 | 2.30 ± 0.15 | |
| $K_{m2}$ (μM) | 449.8 ± 828.4 | | 2223 ± 578.4 | 43.99 ± 23.14 | |

### Mechanism of ion selectivity in K^+^-independent H^+^-PPases

The structure of the ion gate must hold the explanation to ion selectivity, but the current model, that the position of the semi-conserved glutamate defines ion selectivity (6.53 in Na^+^- PPases; 6.57 in H^+^-PPases, does not hold for K^+^-independent H^+^-PPases (see Introduction). The 2mF_o_-dF_c_ map quality at the ion gate of subunit A was good and all residues except for K^16.50^ are defined reasonably well, including the semi-conserved glutamate (Figure 2-figure supplement 2B). We evaluated the different orientations of the rotamer library used by Coot for K^16.50^ to place its side chain. The chosen rotamer does not clash and shows the best hydrogen bonding, making it superior to all other options (Supplementary table 7). In this conformation, it is positioned as in *Vr*PPase:Mg_5_IDP (Figure 5A-B), coordinated to D^6.50^, S^6.54^ and N^16.46^, consistent with *Pa*PPase pumping H^+^. All other side chain orientations and interactions at the ion gate of *Pa*PPase:Mg_5_IDP resembled the *Vr*PPase:Mg_5_IDP structure, despite the shift of the semi-conserved glutamate (E^6.53^ in *Pa*PPase, E^6.57^ in *Vr*PPase)(Figure 5B). This is also consistent with the biology as *both* proteins, *Pa*PPase and *Vr*PPase, are H^+^- PPases – but in this case it is not clear what could reorient E^6.53^ in *Pa*PPase so that it forms the same interactions as E^6.57^ in *Vr*PPase, not as E^6.53^ in *Tm*PPase.

**Figure 5:**
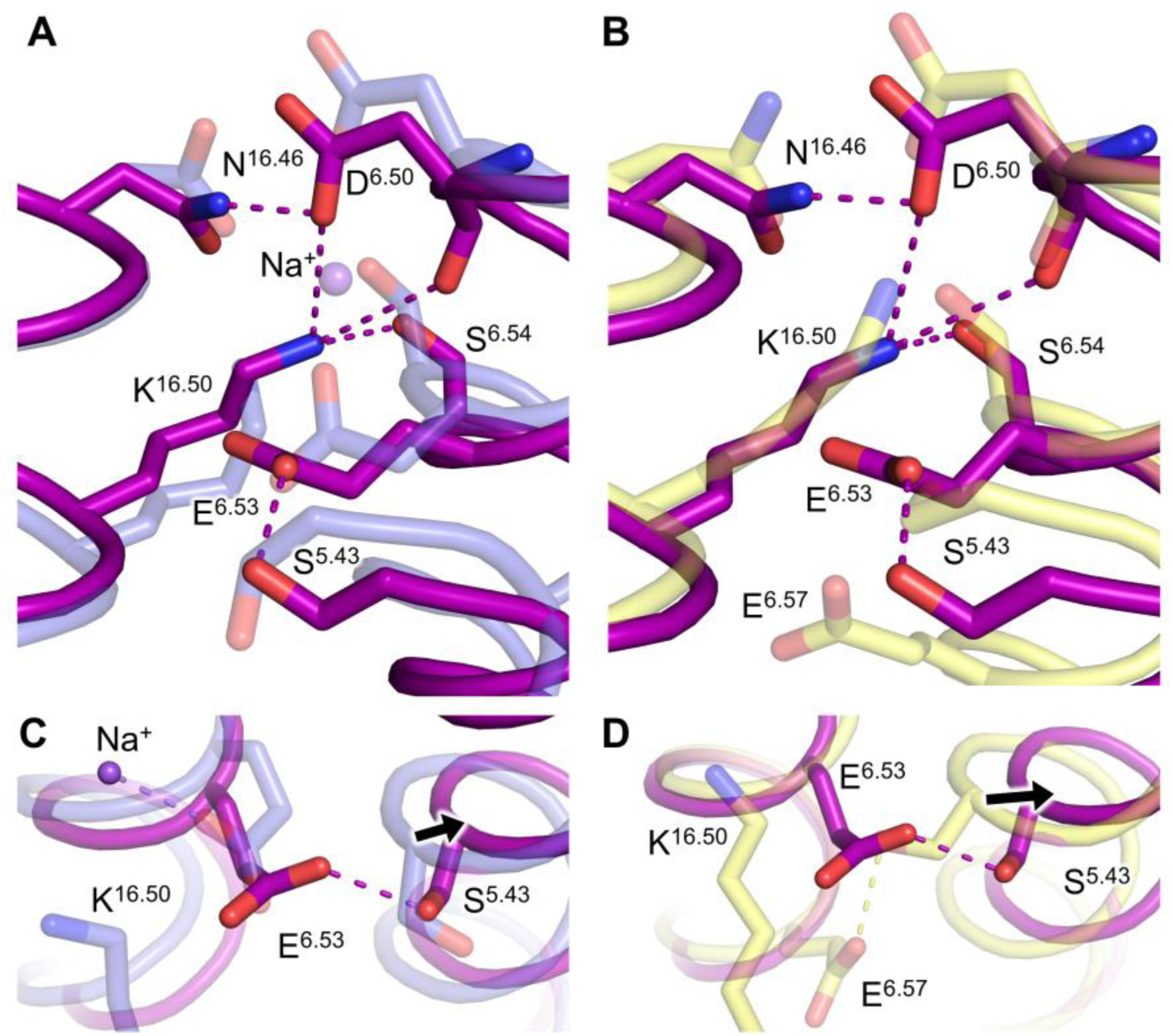
Structural overview of the ion gate in *Pa*PPase:Mg5IDP. (**A**) Comparison of the *Pa*PPase:Mg5IDP (purple) and *Tm*PPase:Mg5IDP (blue) ion gate structures. (**B**) Comparison of the *Pa*PPase:Mg5IDP (purple) and *Vr*PPase:Mg5IDP (yellow) ion gate structures. (**C**) Close-up view and comparison of the semi-conserved glutamate (E^6.53/57^) orientation and helix 5 conformation in *Pa*PPase:Mg5IDP (purple) and *Tm*PPase:Mg5IDP (blue). (**D**) Close-up view and comparison of the semi-conserved glutamate (E^6.53/57^) orientation and helix 5 conformation in *Pa*PPase:Mg5IDP (purple) and *Vr*PPase:Mg5IDP (yellow). Dashed lines shown the coordination of key residues involved in ion selectivity such as S^5.43^, E^6.53^ and K^16.50^. Major structural changes are indicated by black arrows.

In *Pa*PPase:Mg_5_IDP, helix 5 straightened and moved out of the protein core at the ion gate by about 2 Å compared with other IDP-bound M-PPases (Figure 5C-D). This allows S^5.43^ to hydrogen bond to E^6.53^, as occurs with E^6.57^ in *Vr*PPase:Mg_5_IDP (Figure 1D). In *Tm*PPase, helix 5 is closer to helix 6 and helix 16, forcing position 6.53 to point away from S^5.43^, thereby contributing to the formation of a Na^+^-binding site (*Tm*PPase:Mg_5_IDP)(Figure 5C).

### An ion-binding site at the dimer interface

The dimer interface of *Pa*PPase is formed by helices 10, 13 and 15 and somewhat different to other M-PPases (Figure 5-figure supplement 1A). Usually, non-polar amino acids are conserved at position 10.44 (97.6% conserved) and 15.49 (95.2% conserved). In *Pa*PPase these are tyrosine and arginine, respectively (Figure 5-figure supplement 1B). The additional hydrogen-bonding potential and positive charge leads to the formation of an anion binding site. We modelled SO_4_ ^2-^ from the crystallisation solution to mediate the inter-subunit communication of Y^10.44^, Y^13.44^, and R^15.40^ in *Pa*PPase:Mg_5_IDP, but this may be P_i_ under physiological conditions.

### Direct observation of asymmetric PP_i_ binding in *Tm*PPase

Structural information about inter-subunit communication and functional asymmetry is essential to resolve unanswered questions about variable ion pumping selectivity and, potentially, energy coupling in M-PPases (see Introduction). Unfortunately, structural data on symmetric, inhibited enzyme can only provide limited insight. We therefore decided to study the K^+^-dependent Na^+^-PPase from *T. maritima* (*Tm*PPase) using a time-resolved cryo-trapping approach to be able to map asymmetric enzyme states.

The catalytic turnover (*k*_cat_) of purified *Tm*PPase that was crystallised (Figure 6-figure supplement 1A) in conditions suitable for time-resolved studies (*i.e.* no inhibitors) was 282-fold lower (*k*_cat_: 0.16 ±0.05 s^-1^ at 20 °C) compared to ideal reaction conditions (*k*_cat_: 45.13 ±3.59 s^-1^ at 71° C)(Figure 6-figure supplement 1B, Supplementary table 8), of which about a factor of 32-234 may be ascribed to the change in temperature assuming a Q_10_ of 2-3 as for most biological systems (Blehrádek, 1926). The protein was thus active and, importantly, the substrate turnover sufficiently slow for a manual single-crystal time-resolved cryo-trapping approach in which the reaction was initiated by soaking-in activating Na^+^ (see Methods).

**Figure 6:**
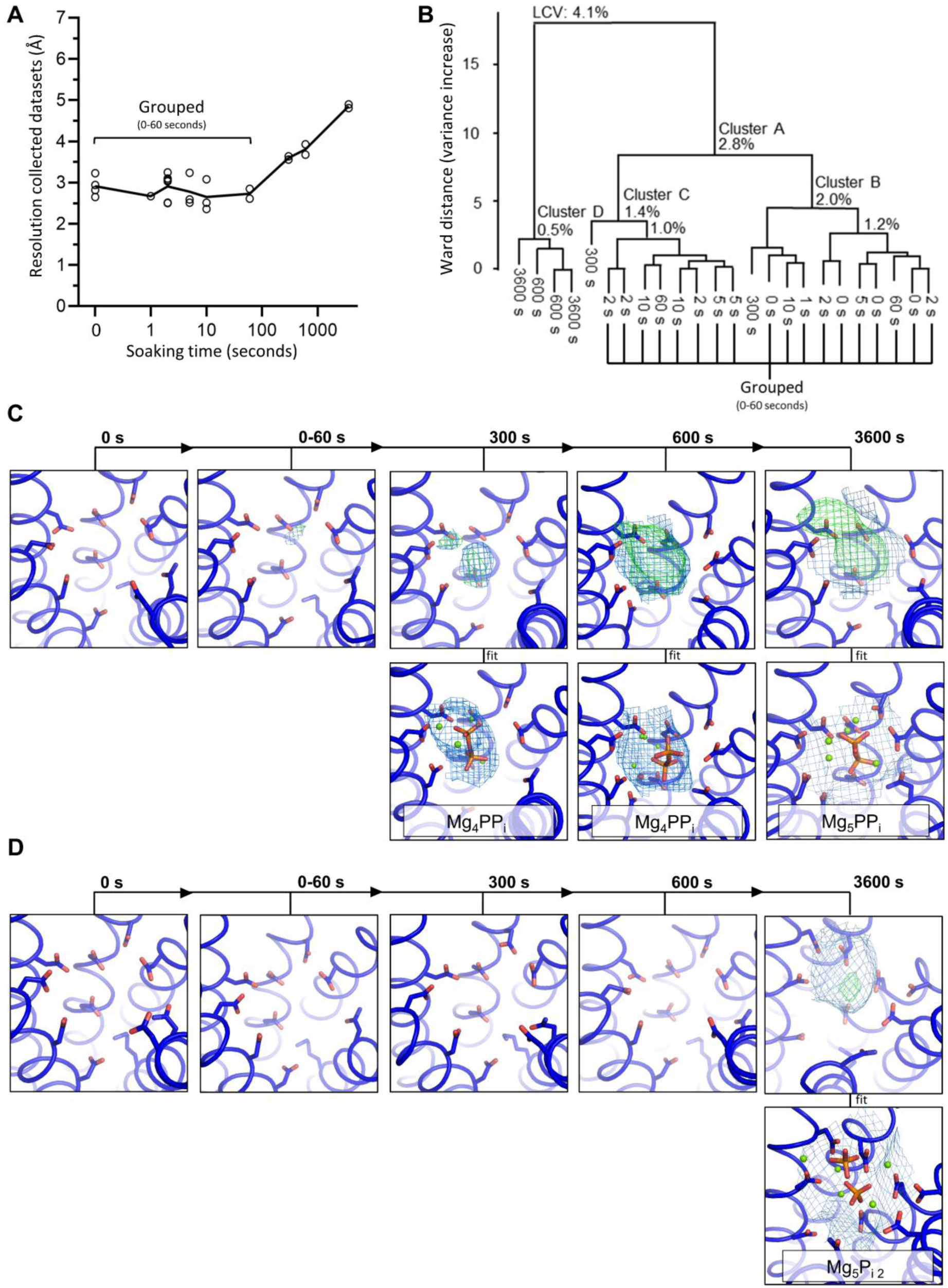
Characterisation of time-resolved *Tm*PPase datasets. (**A**) Diffraction quality at different time-points. Each collected dataset is represented by a circle with the diffraction in the best direction plotted. The mean resolutions of *each* time point are connected by a black line. (**B**) Dendrogram of BLEND analysis to identify isomorphous time-resolved datasets. Nodes of the four biggest cluster are labelled with the linear cell variability (LCV). (**C-D**) *Tm*PPase active site of subunit A (**C**) and subunit B (**D**) at different time-points with 2mFo-dFc density (blue) and mFo-dFc density (red/green) for ligand shown. If not stated otherwise, 2mFo-dFc density is contoured at 1 σ and the mFo-dFc density is contoured at 3 σ.

The collected time-resolved *Tm*PPase datasets (t=0-3600 seconds) were severely anisotropic (Supplementary table 9). Non-soaked reference crystals (t=0 seconds) yielded a structure with a resolution of 2.65 Å along h, 3.32 Å along k and 3.79 Å along l at best (Supplementary table 9). To avoid bias, the *Tm*PPase structures were solved by molecular replacement using the *Tm*PPase:CaMg resting state structure (PDB: 4AV3) as a search model with hetero atoms removed. The 0-seconds structure has one homodimer molecule per asymmetric unit, refined to an R_work_/R_free_ of 23.8/27.4% and was very similar to the inhibited *Tm*PPase:CaMg structure (r.m.s.d./C_α_: 0.41 Å). To check for asymmetry, subunits A and B were refined individually, but remained identical at this resolution, with an r.m.s.d./C_α_ of 0.21 Å. Despite the presence of 0.4 mM PP_i_ in the crystallisation condition, it was not located at the active site, nor did it bind anywhere else.

The diffraction quality of crystals that were soaked in Na^+^-trigger solution for up to 60 seconds was similar to non-soaked reference crystals. Interestingly, the diffraction quality declined abruptly in datasets collected at >60 seconds (Figure 6A). The diffraction limits for the 300-, 600- and 3600-seconds time-points were 3.77 Å, 3.84 Å and 4.53 Å in the best direction, respectively. By trying different combinations of datasets, it became clear that all of the data from 0-60 seconds could be combined into one, yielding a new structure of *Tm*PPase in the resting state (*Tm*PPase:Mg_2_) that is essentially identical to the *Tm*PPase:CaMg (r.m.s.d./C_α_: 0.29 Å) and 0-seconds structure (r.m.s.d./C_α_: 0.33 Å). This led to improved data quality parameters including resolution and completeness, while R_pim_ remained stable and within the generally accepted limit of ∼5% (Supplementary table 9). The structure was solved at 2.54 Å along h, 2.95Å along k and 3.38 Å along l with improved B-factors (108.65 Å^2^ to 70.28 Å^2^) and R_work_/R_free_ values (21.95/23.61%) compared to the 0-seconds structure (23.81/27.40%). The improvement in data quality parameters also translated into better electron density maps, so we could model most side chains and build additional key loop regions, for example loop_5-6_. The changes in diffraction quality also aligned with BLEND analysis (Aller et al., 2016) of the linear cell variability (Figure 6B) in which the 0-60-seconds datasets cluster well (with linear cell variabilities (LCV) of <2.8%). We excluded the 300- seconds data from the combined data set due to the loss in resolution at >60 seconds even though it clustered with a LCV of <2.8%.

Upon calculating difference Fourier maps for the later time points using the 0-60 seconds structure for phases, we were able to observe, for the first time, significant asymmetry in an active M-PPase, corresponding to the first steps in the catalytic cycle. There is positive mF_o_-dF_c_ density at 3 σ in subunit A but none in subunit B at 300 and 600 seconds; and the position of that positive density changes between these two time points (Figure 6C-D).

We then combined all datasets obtained at the same time-point, which likewise improved electron density maps and data quality parameters (Supplementary table 9). At 300 seconds, the density of combined datasets is best fit by full-occupancy PP_i_ in subunit A (Figure 6C). Although placing lower-occupancy PP_i_ leads to lower B-factors (Supplementary table 10), positive mF_o_-dF_c_ density remains after refinement, and R_work_ and R_free_ are higher. At this time- point, PP_i_ has not arrived yet in its final binding pose and is tilted when compared to IDP binding (r.m.s.d: 1.5 Å) in *Tm*PPase:Mg_5_IDP (Figure 7A). The phosphor atom of the leaving group phosphate is displaced by 1.4 Å and oxygen atoms by up to 1.9 Å (Figure 7B). This is consistent with the helix orientations remaining highly symmetrical between both subunits (r.m.s.d./C_α_: 0.21 Å), and very similar to the resting-state *Tm*PPase:CaMg (r.m.s.d./C_α_: 0.35 Å) and the 0-60-seconds structures (r.m.s.d./C_α_: 0.13 Å). After 600 seconds, PP_i_ is bound at the canonical position for hydrolysis (Figure 7C). Consequently, loop_5-6_ was modelled to seal the active site and helix 12 had shifted downward (Figure 7D) as seen in *Tm*PPase:IDP, *Pa*PPase:IDP and *Vr*PPase:IDP structures, all while subunit B remained in the resting state (r.m.s.d./C_α_ subunit A *vs.* B: 0.79Å).

**Figure 7:**
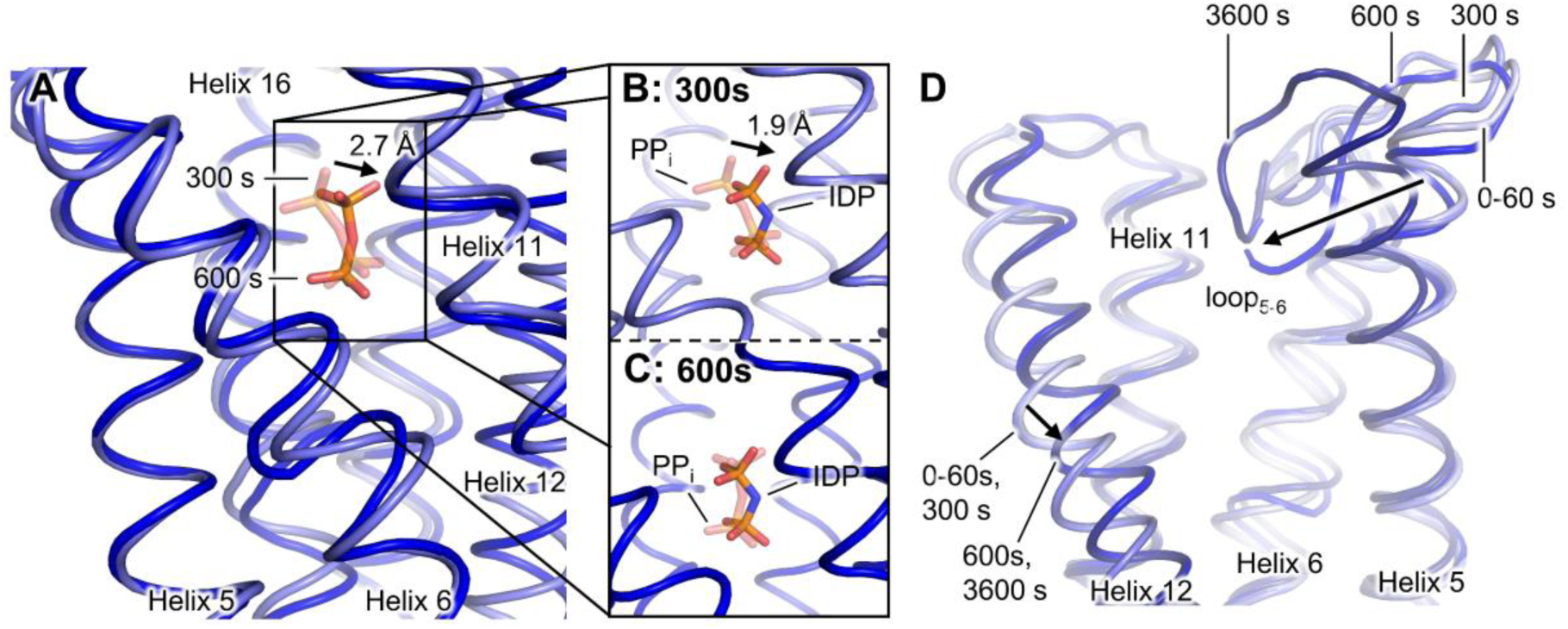
Structural comparison of the 0-60-seconds, 300-seconds, 600-seconds and 3600-seconds time- resolved *Tm*PPase structures. (**A**) Comparison of the PPi binding position in the 300-seconds (light blue) and 600-seconds (blue) structure of *Tm*PPase. (**B**) Comparison of the PPi (transparent) binding position in the 300-seconds structure with the IDP binding position in *Tm*PPase:Mg4Pi2. (**C**) Comparison of the PPi (transparent) binding position in the 600-seconds structure with the IDP binding position in *Tm*PPase:Mg5IDP. (**D**) Helix reorientation and loop5-6 closure upon substrate binding in the 0-60-seconds, 300-seconds, 600-seconds and 3600-seconds structure (labelled and coloured in different shades of blue from light to dark). Black arrows highlight major structural changes.

In the 3600-seconds structure, both active sites appear to be occupied (Figure 6C-D), which demonstrates unrestricted access and shows that asymmetric binding is not a crystallographic artefact. The loss of resolution upon binding makes analysis very difficult indeed, but the mF_o_-dF_c_ maps (Figure 6C-D) and B-factor distributions (Supplementary table 10) are not inconsistent with the idea that, at this point, one subunit binds PP_i_ and the other 2 P_i_. Intriguingly, the density is fit best by PP_i_ still present in subunit A and 2 P_i_ bound in subunit B. In accordance with this, the main chain electron density is fit best when subunit A is modelled in the closed state as seen in IDP-bound structures (r.m.s.d./C_α_ to *Tm*PPase:Mg_5_IDP: 0.49 Å), and subunit B modelled in the open state as seen in resting- or product-bound structures (r.m.s.d./C_α_ to *Tm*PPase:CaMg: 0.45 Å).

These structures are completely consistent with the half-of-the-sites reactivity seen in kinetic assays (Anashkin et al., 2021; Artukka et al., 2018; Vidilaseris et al., 2019); they provide snapshots of a structural binding pathway, and support a new comprehensive kinetic model of catalysis (see Discussion).

There were no significant changes in other regions of the protein, but the semi-conserved glutamate E^6.53^ appears to be flexible in all the structures as indicated by the absence of 2mF_o_-dF_c_ density or negative density when modelled as seen in *Tm*PPase:CaMg (Figure 6- figure supplement 2). All other side chain orientations at the ion gate were well defined in the 2mF_o_-dF_c_ map up to the 300-seconds time-point. Typically, the semi-conserved glutamate is well ordered and side chain density visible at much lower resolutions (see *Pa*PPase structure, Figure 2-figure supplement 2B). Its flexibility in low Na^+^ conditions may have implications for the ion pumping selectivity at sub-physiological Na^+^ concentrations in K^+^-dependent Na^+^- PPases (see Discussion).

### Energy coupling in M-PPases

We further used SURFE^2^R N1 to study energy-coupling of PP_i_ hydrolysis and Na^+^ pumping in *Tm*PPase. Currents were generated when ions crossed the membrane, so the measured current is the sum of the currents from all active proteins on the sensor. A positive signal of 0.5±0.05 nA was detected after addition of 100 μM substrate (K_4_PPi) and reached its maximum within ∼0.1 seconds, in line with the dead time of the machine under the conditions used (Figure 8). This is at least 12-25 times faster than the turnover rates at 20 °C under similar lipidated conditions (Figure 6-figure supplement 1C, Supplementary table 8). We thus propose that electrometric signals correspond to a single pumping event and that the overall turn-over is rate limited by PP_I_ hydrolysis or phosphate release. Indeed, we observed that to fully recover the signal on the same sensor, a waiting time of several minutes was required, in line with the proposition that hydrolysis or phosphate release is slow. When the substrate was replaced by 50 μM of the non-hydrolyzable analogue IDP, the signal was reduced by half (Figure 8). In the presence of 200 μM K_2_HPO_4_ as a negative control, the current was less 0.02 nA. The signal decayed within about 2 seconds, corresponding to *Tm*PPase entering a state that temporarily could no longer pump Na^+^ at a sufficient rate to generate a signal. The current decay curves (Figure 8 B-C) were well fit by a single exponential with similar decay rates (*k)* for PP_i_ (6.0±0.4 s^-1^) and IDP (9.6±1.1 s^-1^) (Table 3). Overall, these measurements are consistent with *Tm*PPase pumping Na^+^ upon addition of either PP_i_ or IDP (see Discussion), with PP_i_ generating two pumped Na^+^ and IDP one.

**Table 3:**
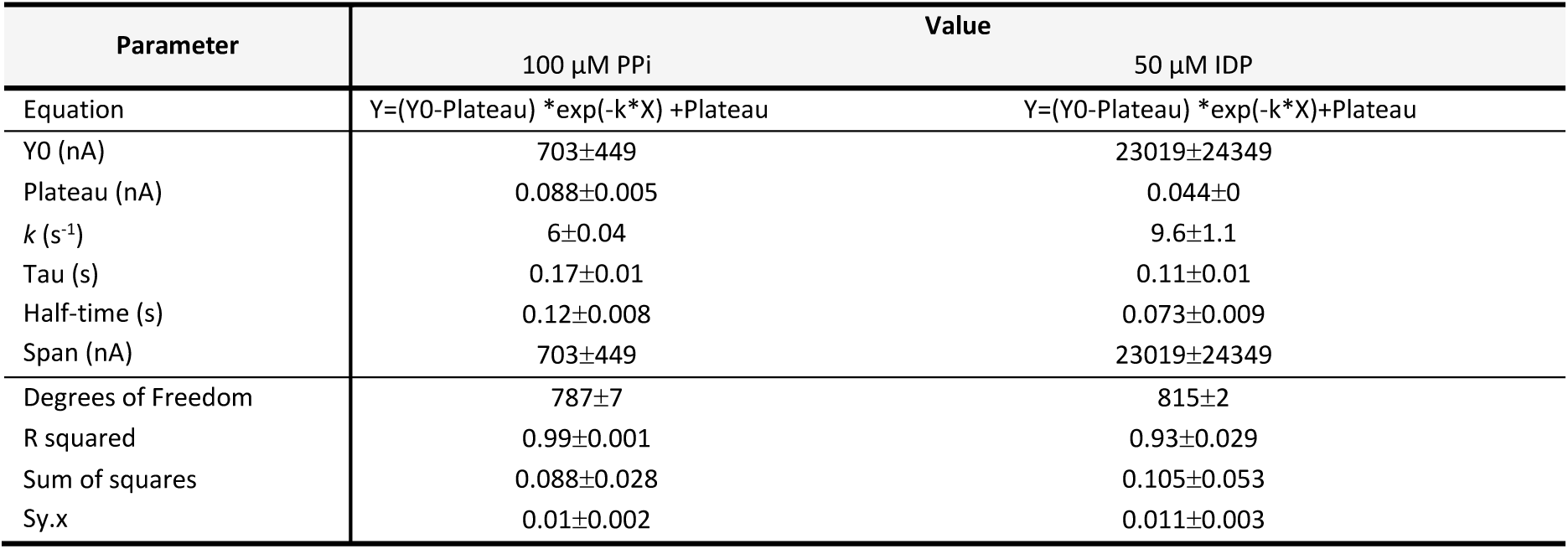
Current decay parameters for Na^+^ pumping of TmPPase.

**Figure 8:**
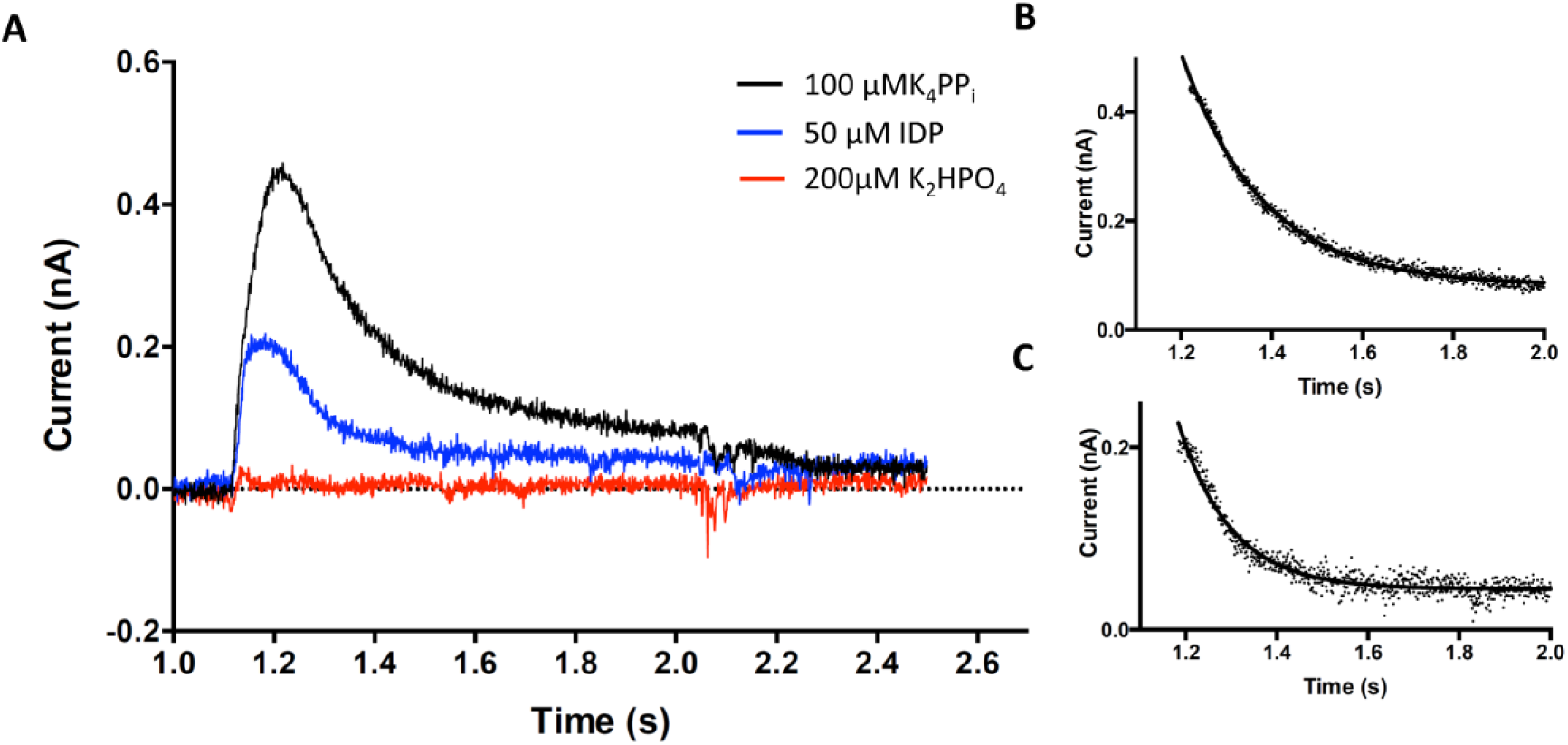
Transient currents of *Tm*PPase Na^+^ pumping. (**A**) Triggered by 100 µM of K4PPi, 50 µM of IDP and 200 µM of K2HPO4. (**B**) Current exponential decay fit curve of PPi (1.2 to 2 seconds). (**C**) Current exponential decay fit curve of IDP (1.2 to 2 seconds).

## Discussion

Our study and recent work has clearly indicated that M-PPases show anti-cooperative behaviour: productive substrate binding cannot happen in both sites at the same time (**Table 2**) (Anashkin et al., 2021; Artukka et al., 2018; Vidilaseris et al., 2019). Artukka and co-workers proposed a model where binding in subunit A converts subunit B into a conformation that prevents substrate binding - even though all published structures of M-PPases with IDP show symmetrical binding to both subunits (Artukka et al., 2018). Vidilaseris *et al*. (Vidilaseris et al., 2019), who identified an allosteric inhibitor of *Tm*PPase, suggest that motions of loops near the exit channel lead to asymmetry and play a role in intra-subunit communication. In their structure, these changes create a binding site for the allosteric inhibitor ATC in subunit A and prevent full motion in subunit B.

It is from this background we endeavour to synthesise a comprehensive model of M-PPase catalysis. An ideal model would explain (a) half-of-the-sites reactivity; (b) energy-coupling of hydrolysis and ion pumping; (c) varying ion selectivity; and (d) how certain pumps can pump both Na^+^ and H^+^ using the same machinery. It is our contention that one, unified model that puts intersubunit communication at the *heart* of the catalytic cycle explains all of these.

### Half-of-the-sites reactivity

We start with a structural explanation of half-of-the-sites reactivity in the context of our new data. *Pa*PPase, and K^+^-independent M-PPases in general, have additional hydrogen bonding between helix 11 and 12 through T^12.49^-D^11.50^ (*Plasmodium spp.* S^11.50^), that when lost in T^12.49^A mutants, abolishes activity. Moreover, the K^12.46^A mutation eliminates half-of-the-sites reactivity in *Pa*PPase that can only be restored by addition of K^+^: positive charge around the 12.46 sidechain position and the motion of helix 12 is a key part of communication. This is in line with the model of intra-subunit ion gate - ion gate communication *via* exit channel loops, particularly loop_12-13_, that was proposed by Vidilaseris *et al*. and directly links the inner ring helices such as helix 12 to the dimer interface (Vidilaseris et al., 2019). Intriguingly, the length of the exit channel loops appears to be conserved: if exit channel loop_8-9_ is long (12-18 residues), loop_10-11_ is short (3-9 residues) and *vice versa* (Figure 9-figure supplement 1). A second communication network is from helix 5 to helix 13 and propagated through helix 10 (Figure 3B-C, Figure 3-figure supplement 2).

**Figure 9:**
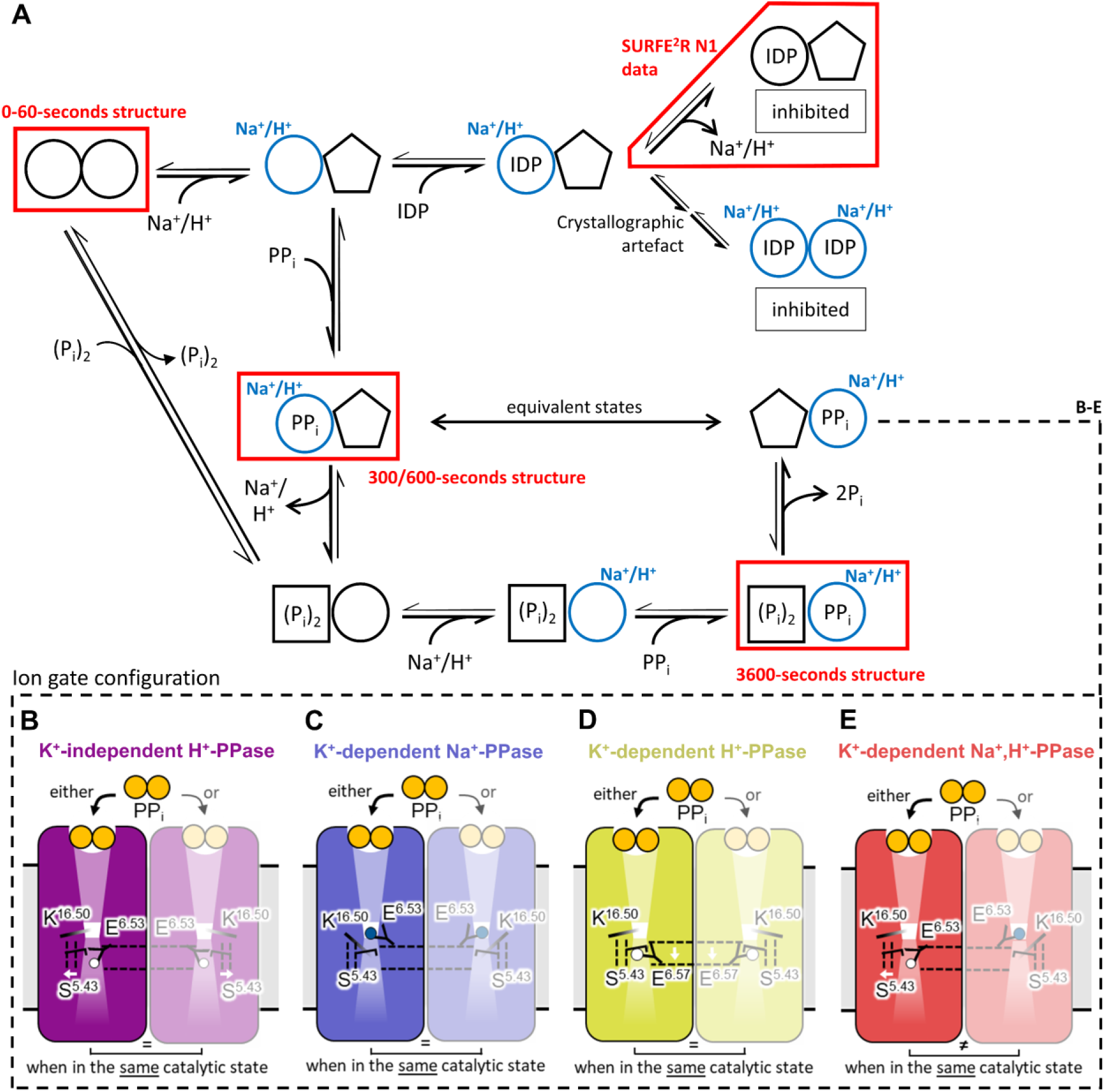
Unified model of M-PPase catalysis. (**A**) Schematic model of functional asymmetry. The active site status is defined by shape, and binding of the pumped ion at the active side indicated by a label and blue colouring. PPi can only bind to one subunit at a time and requires prior binding of the pumped ion at the ion gate. The thermodynamically favoured reaction pathway is indicated by bold arrows. In single-pumping M-PPases, the ion gate conformation of subunit A and subunit B are mirror images of each other (equivalent states – see panel B-D). In dual-pumping M-PPases, the ion gate conformation of subunit A and subunit B are asymmetrical (*i.e.* H^+^- pumping setup *vs.* Na^+^-pumping setup – see panel E). Enyzme states mapped by time-resolved crystallography or investigated in electromagnetic studies are highlighted by red boxes. (**B-E**) Schematic model of ion selectivity with the orientation of key residues K^16.50^ (|) E^6.53/57^ (Y) and S^5.43^ (˥) shown. The ion gate set up is shown with both subunits in the same catalytic state for comparison. Under physiological conditions, subunit A and subunit B are never in the same catalytic state according to the half-of-the-sites reactivity model. The position of helix 5 and the semi-conserved glutamate are indicated by dashed vertical and horizontal lines, respectively. Conformational changes or residue repositioning are indicated by a white arrow. (**B**) E^6.53^ protonation (white sphere) and interaction with S^5.43^, made possible by the outward movement of helix 5. K^16.50^ destroys the Na^+^ binding site. (**C**) Na^+^ (purple sphere) binding to the ion gate and interaction with E^6.53^, which faces away from S^5.43^ when helix 5 is close to avoid clashing. (**D**) E^6.57^ protonation as a results of its shift one helix turn down, which creates sufficient space for the S^5.43^-E^6.53^ interaction. (**E**) Proposed functional asymmetry at the ion gate of in K^+^-dependent Na^+^,H^+^-PPases, explaining Na^+^ binding and E^6.53^ protonation in the respective subunit. Ion selectivity depends on helix 5 orientation and E^6.53^-S^5.43^ interaction. Subunit A is similar to the residue and helix orientation shown in panel B, whereas subunit B is similar to the residue and helix orientations of panel C.

In our model, binding of the ion in subunit A followed by substrate binding (**Figure 9A**: 300-seconds structure) leads to the downward motion of helix 12 and straightening and tight winding of helix 5, and is propagated into subunit B *via* helices 5,12, 13 and the exit channel loops. Implicit in this is the idea that (ion) binding and pumping in subunit B can only happen once subunit A has reset (Figure 9A).

Our evidence for this new model comes from the time-resolved cryo-trapped structures (Figure 6), kinetic data (Table 2) (Anashkin et al., 2021; Artukka et al., 2018; Vidilaseris et al., 2019) and the SURFE^2^R N1 data (Table 3). First, catalysis as measured by phosphate production has a *k*_cat_ of 0.2-0.4 s^-1^ at 20 °C (Supplementary table 8) but maximal signal in the SURFE^2^R N1 is reached in the dead time of the machine (∼0.1 seconds), and the height of the peak for PP_i_ is about twice that of IDP (Figure 8). Consequently, the rate of pumping as measured by the SURFE^2^R N1 is at least 12-25 times that of PP_i_ hydrolysis/phosphate release. The decay constants (protein unable to pump) for the two are similar (6±0.4 s^-1^ for PP_i_ and 9.6±1.1 s^-1^ for IDP), suggesting that they correspond to a similar event. We thus hypothesize that PP_i_ hydrolysis/phosphate release from the liposomes at 20 °C is rate limiting and extremely slow for the thermophilic *Tm*PPase. The logical explanation is that PP_i_ can bind and pump two Na^+^ (one from each subunit, with a complete catalytic cycle happening at least in subunit A), while IDP only binds to one and pumps one (Figure 9A), as ion and substrate binding to subunit B cannot take place until hydrolysis happens in subunit A.

Second, the asymmetric time-resolved *Tm*PPase structures indicate a highly-ordered reaction mechanism. Ion binding is the first event, as there is no evidence of PP_i_ in any of the structures before 60 seconds and the ion gate E^6.53^ is disordered: Na^+^ binding precedes substrate binding (Figure 6). PP_i_ appears to bind in subunit A in two modes: a distal mode where the electrophilic phosphate is tilted out of its final binding position (t=300 seconds), followed by a hydrolysis-competent mode (t=600 seconds). However, there is no evidence up to 600 seconds of reaction initiation of any binding in subunit B, which remains in the resting state. Even though the resolution is poor, the 3600-seconds structure seems to contain a PP_i_ in subunit A but (P_i_)_2_ in subunit B, suggesting at least one full turnover (Figure 9A) may have occurred.

### Ion selectivity

We can extend the model presented above to explain ion selectivity. The current model (Li et al., 2016) posits that M-PPases with E^6.53^ are Na^+^-pumps, but ones with E^6.57^ are H^+^-pumps (Nordbo et al., 2016) - but both Na^+^,H^+^-PPases and K^+^-independent H^+^-PPases have E^6.53^ (**Table 1**). However, we propose that ion selectivity depends on the interaction between the semi-conserved glutamate with a highly conserved serine on helix 5, S^5.43^ (pairwise identity: 90.6%). This interaction, propagated through the dimer interface as described above, mediates helix 5 orientation and ion selectivity/binding. The ion gate configuration is thus directly linked to the orientation of helix 5, which determines whether E^6.53^ points towards S^5.43^, accommodating a proton, or away from S^5.43^, forming a Na^+^-binding site with D^6.50^, S^6.54^ and D/N^16.46^ (Video 1). In what follows, we show that this mechanism is valid for all currently known M-PPase subclasses (Figure 9B-E).

In K^+^-independent H^+^-PPases, protonated E^6.53^ is held in place by S^5.43^ (Figure 9B), while in Na^+^- PPases, the bent helix 5 forces E^6.53^ to point away from S^5.43^ to avoid clashing, creating a Na^+^-binding site (Figure 9C). Our *Tm*PPase structure obtained in the absence of Na^+^ (0-60-seconds structure) suggests that in the absence of Na^+^, E^6.53^ is not bound by K^16.50^, allowing it to interact with S^5.43^. This could thus explain why K^+^-dependent Na^+^-PPases can pump protons at Na^+^ concentrations <5 mM (Luoto, Nordbo, et al., 2013). In K^+^-dependent H^+^- PPases, E^6.57^ uncouples ion selectivity from the orientation of helix 5 as it can be coordinated by S^5.43^ independent of helix 5 geometry (Figure 9D). The unusual *Flavobacterium johnsoniae* (Luoto et al., 2011) H^+^-PPase, where the semi-conserved glutamate is at position 5.43, is also consistent with our model. The side chain carboxylate of E^5.43^ cannot promote Na^+^-binding about 5 Å away, but can be modelled to occupy a position similar to that of E^6.57^ in *Vr*PPase:Mg_5_IDP, allowing protonation and deprotonation of E^5.43^ (Tsai et al., 2014). In all these structures, ion binding in subunit B would not occur before pumping and hydrolysis in subunit A, but the structure of the ion binding site would be the same.

Finally, to explain ion pumping selectivity in K^+^-dependent Na^+^,H^+^-PPases, we posit that the ion gate configuration flips between the H^+^-PPase (*Pa*PPase:Mg_5_IDP) and Na^+^-PPase (*Tm*PPase:Mg_5_IDP) conformations (this also appears to be possible in K^+^-dependent Na^+^-PPases at low Na^+^ concentrations (Luoto, Nordbo, et al., 2013)). For instance, if subunit A is H^+^-pumping, conformational changes through the helix 5-13 connection are not only required to allow ion binding in subunit B but also to convert it into the Na^+^-binding conformation and *vice versa* (Figure 9E). Seen this way, the K^+^-dependent Na^+^,H^+^-PPases use a special class of the standard M-PPase mechanism, rather than being *sui generis.* We also suggest that *all* M-PPases, not just the nine studied so far (Anashkin et al., 2021; Artukka et al., 2018; Vidilaseris et al., 2019), will show half-of-the-sites reactivity.

Conversely, two recently-published papers from Baykov and co-workers (Baykov et al., 2022; Malinen et al., 2022) continue to posit their “billiard-type” mechanism, where hydrolysis precedes pumping. By using stopped-flow measurements on *Tm*PPase at 40 °C, they demonstrate that hydrolysis is the most likely rate-determining step, which is consistent with our SURFE^2^R N1 data on *Tm*PPase at 20 °C. They propose that this means that hydrolysis occurs at the same time or precedes pumping – *i.e.* the chemical proton released from hydrolysis enters a Grotthus chain and forces the pumped Na^+^ into the exit channel. However, their pre-steady-state rate at 40 °C is 12 s^-1^, consistent with the *k*_cat_ of our steady state kinetics at 20 °C (0.2-0.4 s^-1^) (Supplementary table 8). The pre-steady-state rate at 20 °C is thus likely to about 1-3 s^-1^ (a factor of 2-3 per 10 °C)(Blehrádek, 1926) - *i.e.,* at least 4-10 times slower than the rates observed in the Nanion Surfer, where PP_i_ binding and ion transfer happens within the dead time of the machine (≈0.1 seconds) and the decay following the initial pumping event is of a similar speed (8.8 s^-1^). The simplest explanation, to borrow from Dr. Doolittle (Wikipedia Community, 2022), is thus not a pushmi-pullyu mechanism, but a pullyu-pushmi mechanism: the pumping of the Na^+^ increases the negative charge in the closed active site, causing deprotonation along the Grotthus chain and hydrolysis of the PP_i_. We concur with Baykov and co-workers that, whatever the mechanism, it must the same for all M-PPases as the catalytic machinery is so similar. We also agree that the most likely identity of the charged residue, as identified by their steady-state solvent isotope experiments, is indeed E^6.57^ with a *p*K_a_ of about 7.8. A glutamate in the membrane that has just lost a counter-ion would suit perfectly.

In summary, the only sequence of events that is consistent with all the data presented above, including the pre-steady state experiments by Baykov and co-workers (Malinen et al., 2022), is based on a half-of-the-sites reactivity mechanism, supports a binding-change-type (“pumping-before-hydrolysis”) energy coupling and is as follows (Figure 9A): Binding of ion at the gate in monomer A allows binding of substrate, which prevents binding of ion or substrate at monomer B. Pumping in monomer A leads to hydrolysis in monomer A, which releases monomer B into an ion-binding/substrate binding conformation. Ion-binding in monomer B could then allow release of product in monomer A, followed by substrate binding and pumping in B. The beauty of this mechanism is that it provides a convincing rationale for all of the observed data, in particular explaining half-of-the sites reactivity and dual pumping M-PPases. Other models, positing that two ions can occasionally be pumped in one subunit are, to our mind, not as convincing as no modern experiments on purified proteins have indicated a hydrolysis/transport ratio above 1:1.

Our model provides, for the first time, an overall explanation of ion selectivity and catalysis in *all* M-PPases and makes testable predictions: *e.g.* global conformational changes should occur in the first 0.1-0.2 s at 20 °C. These need to be tested through functional and structural studies, in particular the use of time-resolved, single-molecule techniques to capture further details of mechanism, as well as efforts to capture mechanistic details through molecular dynamics – efforts that are already underway (Holmes et al., 2022).

## Materials and Methods

### Mutagenesis of *Pa*PPase

We used N-terminally RSGH_6_-tagged constructs for full-length *Tm*PPase expression from (Kellosalo et al., 2011) and replaced the open reading frame encoding for *Tm*PPase with a section encoding for full-length *Pa*PPase instead. The *Pa*PPase gene was PCR amplified (Q5® Hot Start High-Fidelity 2X Master Mix from NEB, Frankfurt am Main, Germany) with primers introducing a GG-linker along with a 5’ SalI (TTT TTT GTC GAC ATG CAT CAC CAT CAC CAT CAC GGT GGA AAT ATG ATA AGC TAT GCC TTA CTA GG) and 3’ XbaI (TTT TTT TCT AGA TCA GAA AGG CAA TAG ACC TG) restriction site. The PCR product was inserted into the linearised (SalI, XbaI from NEB) pRS1024 yeast expression vector (Kellosalo et al., 2011). *Pa*PPase variants K^12.46^A (C AAT ACC ACA gca GCC ACT ACT AAG GG, CC GAC GGA GTC CAG TAC A), T^12.49^A (A AAA GCC ACT gct AAG GGA TAT GC, GT GGT ATT GCC GAC GGA G) and the combination of both were generated using the Q5® site-directed mutagenesis kit (NEB) (lower case letters highlight the amino acid change). Template DNA was removed by DpnI (NEB) digestion and the constructs were sequenced to confirm the introduction of point mutations.

### Protein expression and purification

We expressed and purified *Pa*PPase and *Tm*PPase in *Saccharomyces cerevisiae* as described elsewhere (López-Marqués et al., 2005; Strauss et al., 2018). In brief, yeast expression plasmids carrying N-terminally 6xHis-tagged wild-type or variant *Pa/Tm*PPase under control of the constitutively active PMA1 promoter were freshly transformed into the *S. cerevisiae* strain BJ1991 (genotype: *MATα prb1-1122 pep4-3 leu2 trp1 ura3-52 gal2*) and cultivated at 30 °C for 12 hours in 250 mL selective synthetic complete dropout starter cultures lacking leucine (SCD-Leu, in-house). 750 mL of 1.5x SCD-Leu (*Pa*PPase) or 1.5x yeast peptone dextrose (YPD, in-house)(*Tm*PPase) expression culture were inoculated with 250 mL of starter culture for protein expression at 30 °C. Cells were harvested after 8-10 hours from 10 L expression batches by centrifugation (4,000 *x*g, 4 °C, 15 minutes).

Cells were lysed using a bead-beater (Biospec products, Bartlesville, Oklahoma) with 0.2-mm glass beads and membranes were collected by ultracentrifugation (100,000 xg, 4 °C, 1 hour). The membrane pellet was resuspended in 50 mM MES pH 6.5, 20% *v/v* glycerol, 50 mM KCl, 5 mM MgCl_2_, 2 mM dithiothreitol (DTT), 1 mM phenylmethylsulfonyl fluoride (PMSF) and 2 μg/mL pepstatin to a final total protein concentration of ∼7 mg/mL, mixed with solubilisation buffer (50 mM MES-NaOH pH 6.5 20% *v/v* glycerol, 5.34% w/v n-Dodecyl-β-D-Maltoside (DDM),1 mM K_4_PP_i_) at a 3:1 ratio and incubated at 75 °C (“hot-solve”) for 1.5 hours. Protein was then purified by IMAC using nickel-NTA resin (Bio-Rad, Hercules, California). Depending on the protein and downstream experiments, different buffers were used for purification as outlined below. For structural studies of *Pa*PPase, the solubilised membranes were incubated with nickel-NTA (Cytiva, Marlborough, Massachusetts) resin at 40 °C for 1-2 hours and washed with 2 column volumes (CV) 50 mM MES-NaOH pH 6.5, 20% *v/v* glycerol, 5 mM MgCl_2_, 20 mM imidazole, 1 mM DTT and 0.5% *w/v* n-Decyl-β-D-Maltoside (DM) or 0.05% *w/v* DDM prior to elution in 2 CV 50 mM MES-NaOH, pH 6.5, 3.5% *v/v* glycerol, 5 mM MgCl_2_, 400 mM imidazole, 1 mM DTT and 0.5% *w/v* DM or 0.05% DDM *w/v*. *Pa*PPase samples used in functional studies were solubilised in DDM and contained not MES-NaOH but MOPS-TMAOH (Tetramethylammonium hydroxide)(pH 6.5) instead in order to obtain a “Na^+^-free” sample. The purification of *Tm*PPase followed a similar protocol and using 0.5% *w/v* octyl glucose neopentyl glycol (OGNG) with MES-TMAOH (pH 6.5). In addition, the purification buffers contained 50 mM KCl due to the K^+^-dependence of *Tm*PPase. After nickel-NTA purification, all purified proteins were exchanged into elution buffer lacking imidazole using a PD10 desalting column (Cytiva) and concentrated to ∼10 mg mL^-1^. SDS-PAGE and size exclusion chromatography (SEC) using a Superose^®^ 6 Increase 10/300 GL column (Cytiva) and an NGC Quest 10 Plus System (Bio-Rad) showed that both wild-type and variant proteins were pure and monodisperse (Figure 2-figure supplement 1).

### Vapour-diffusion crystallisation of *Pa*PPase and *Tm*PPase

Initial crystallisation trials of wild-type *Pa*PPase were carried out with several commercial sparse matrix screens using protein solubilised in DM and DDM. Commercial sparse matrix crystallisation screens were set up with protein at 10 mg mL^-1^ (1:1 ratio) after pre-incubation with 4 mM Na_4_IDP (imidodiphosphate) salt or CaCl_2_ (1 hour, 4 °C). Any precipitation that formed within the incubation period was removed by centrifugation at 10,000 xg for 10 minutes prior to setting up crystallisation trials. The best crystals were obtained in the presence of 2 mM IDP in 30-33% *v/v* PEG 400, 0.1 M MES pH 6.5, 0.05 M LiSO_4_, and 0.05 M NaCl at 20 °C with protein solubilised in DM. The crystals were manually harvested at 20 °C. X-ray diffraction was improved by keeping the harvested crystal in the loop for 10 seconds prior to flash cooling in liquid nitrogen, which effectively led to crystal dehydration.

Initial vapour diffusion crystallisation trials of wild-type *Tm*PPase were based on a published crystallisation condition (36% *v/v* PEG 400, 100 mM Tris-HCl pH 8.5, 100 mM MgCl_2_, 100 mM NaCl, 2 mM DTT) (Li et al., 2016) that was further optimised for time-resolved experiments (*i.e.* to contain no inhibitors). *Tm*PPase in OGNG was set up at 10 mg mL^-1^ after pre-incubation with 0.4-4.0 mM K_4_PP_i_ (1h, 4 °C) (instead of Na_4_IDP) and all crystallisation buffers had NaCl replaced with KCl. The best crystals formed in 24-26% *v/v* PEG 400, 50-60 mM Tris-HCl pH 8.5, 2-3 mM MgCl_2_, 175 mM KCl, 2 mM DTT and 0.4 mM K_4_PP_i_ (1:1 ratio) at 20 °C.

### *Pa*PPase data collection, structure solution and refinement

*Pa*PPase crystals were sent to several beamlines including I04 and I24 at the Diamond Light Source (DLS) and ID23-1 and MASSIF-1 at the European Synchrotron Radiation Facility (ESRF) for data collection at 100 K. Collected datasets were processed in XDS (Kabsch, 2010) and the structure was solved by molecular replacement in Phaser (McCoy et al., 2007) using a homology search model based on the 3.5 Å structure of *Tm*PPase:Mg_5_IDP (protein data bank (PDB) ID: 5LZQ) with loop regions removed. The crystals were extremely radiation sensitive, so a complete data set could not be collected on any of them. Consequently, the first few hundred images of eight datasets (3.84-4.35 Å) with positive density for Mg_5_IDP in the active site, less than 2% deviation in unit cell parameters and identical spacegroup (P2_1_) were combined in XDS using XSCALE (Kabsch, 2010). The combined dataset was submitted to the STARANISO webserver (Tickle et al., 2018) prior to molecular replacement as described above (Tickle et al., 2018). Several rounds of refinement using phenix.refine (Liebschner et al., 2019) and manual modelling in Coot (Emsley et al., 2010) were carried out. After an initial round of rigid-body refinement with grouped B-factors, tight restraints were applied to maintain a realistic geometry (torsion angle non-crystallographic symmetry (NCS), secondary structure, and reference structure (PDB: 4A01) restraints). In the last rounds of refinement and Translation-Libration-Screw-rotation (TLS) was enabled and restraints were released except for torsion angle NCS restraints, which were retained to prevent overfitting.

### Time-resolved cryo-trapping X-ray crystallography and structure solution

Time-resolved cryo-trapping crystallography experiments on *Tm*PPase were conducted by manual soaking of vapour diffusion crystals grown in the absence of Na^+^ but in the presence of PP_i_ in a Na^+^-containing trigger solution (60 mM Tris-HCl pH 8.0, 26% *v/v* PEG400, 175 mM KCl, 2.4 mM MgCl_2_, 2 mM K_4_PP_i_, 20 mM NaCl) to initiate the enzymatic reaction *in crystallo*. The reaction was stopped by flash cooling in liquid nitrogen after different soaking times (t=0 [no Na^+^ applied], 1, 2, 5, 10, 60, 300, 600, 3600 seconds) that were selected based on the *k*_cat_ of *Tm*PPase under conditions similar to the crystallisation conditions (Supplementary table 8). Crystallisation wells were re-sealed if the soaking time exceeded 60 seconds to minimise evaporation. Up to five crystals were used for each timepoint.

Diffraction data were collected at 100 K at beamline P14-I at the Deutsches Elektronen Synchrotron (DESY) and the data processed in XDS (Kabsch, 2010) or Xia2/DIALS (Winter et al., 2018). This was followed by anisotropy correction using the STARANISO webserver (Tickle et al., 2018) and molecular replacement in Phaser (McCoy et al., 2007) using the *Tm*PPase:CaMg (PDB: 4AV3) structure without heteroatoms as a search model. The similarity between unit cells of the collected datasets was analysed in BLEND (Foadi et al., 2013) and datasets of the same or different time-points (t=0-60 s) were combined if the linear cell variation was below 3% and the space group and active site status (occupied *versus* not-occupied) were identical.

The single best non-activated structure (reference) and the grouped t=0-60 seconds structure (subset of cluster A) were subject to several rounds of refinement using phenix.refine (Liebschner et al., 2019) and manual modelling in Coot (Emsley et al., 2010). After an initial round of rigid-body refinement with grouped B-factors, torsion angle NCS restraints were applied to further reduce the number of parameters in refinement alongside optimised X-ray/B-factor and X-ray/stereochemistry weighting by phenix. In the final refinement rounds, TLS was applied as well. The 0-60-seconds structure of *Tm*PPase was then used as a search model for molecular replacement of combined datasets that were collected at longer delays after Na^+^-activation. Refinement of these data followed a similar protocol, but the 300-seconds dataset was limited to a single round of 5 refinement cycles, which was sufficient to check for changes of the overall helix geometry at the active site or ion gate. Additional secondary structure restraints were applied in the refinement of the low-resolution t=600 seconds and t=3600 seconds *Tm*PPase structures to maintain realistic geometry.

### Structure analysis

Geneious R11 was used to search the UniProtKB/Swiss-Prot database with blastp (Altschul et al., 1990) for similar sequences to *Pa*PPase and the results were aligned using the Geneious global alignment tool with free end gaps to determine residue conservation and sequence identity. Structure alignments and the r.m.s.d. calculations were done in PyMol 2.2.3 (Schrödinger, LLC, n.d.). The standard deviation was stated when multiple structures were compared by their r.m.s.d.. The solvent accessible surface areas and volumes were determined using HOLLOW with a 1.4-1.5 Å interior probe size (Ho & Gruswitz, 2008). Inter-atom difference distance matrices (DiDiMa) of Cα atoms were generated by the Bio3D R-package for structural bioinformatics (Grant et al., 2006). Hydrogen bonding patterns were analysed in HBplus using default settings (McDonald & Thornton, 1994). The local (residue by residue) helix curvature analysis was done considering blocks of 4 residues using the Bendix plugin of the Visual Molecular Dynamics suite (Dahl et al., 2012), whereas the global (helix by helix) curvature analysis was done using the HELANAL-Plus webserver(Kumar & Bansal, 2012).

### Fixed-time P_i_-release assay for activity measurements under crystallisation conditions

The hydrolytic activity of purified *Tm*PPase for time-resolved structural studies was assessed by using the molybdenum blue reaction method with relipidated (12 mg mL^-1^ L-α-lecithin) protein in DDM:OGNG mixed micelles as previously described (Baykov et al., 2021; Strauss et al., 2018). The reaction buffers were matched to the crystallisation conditions in order to estimate time scales of substrate turnover *in crystallo*. The concentration of MgCl_2_ and K_4_PP_i_ required to maintain 5 mM free Mg^2+^ at pH 8.0 was approximated as described by Baykov and co-workers (Baykov et al., 1993). As reference, a routine reaction was done in 60 mM Tris-HCl pH 8.0, 5 mM free Mg^2+^, 100 mM KCl, and 20 mM NaCl at 71 °C for 5 min. Subsequent reactions at 20 °C were incubated for 240 minutes instead as this produces detectable reaction product in a linear range. The activity of protein in the optimised vapour diffusion crystallisation condition (60 mM Tris-HCl pH 8.0, 100 mM KCl, 3 mM MgCl_2_, 175 mM KCl, 26% *v/v* PEG400, 400 µM K_4_PP_i_) was tested upon reaction initiation with 20 mM NaCl with and without relipidated sample. The standard error of the mean (SEM) was obtained from three technical repeats.

### Continuous-flow P_i_-release assay

Kinetic experiments for wild-type and variant *Pa*PPase were done using phosphate analyzer (Baykov et al., 2021) with relipidated (12 mg mL^-1^ L-α-lecithin) protein in DDM micelles. The reaction mixture of 40 mL contained 50 mM MOPS-TMAOH buffer (pH 7.2) and varying concentrations of free Mg^2+^(added as MgCl_2_) and TMA_4_PP_i_ in ratios that gave the desired concentration of Mg_2_PP_i_ as substrate (Baykov et al., 1993). Reactions were initiated by protein (at low substrate concentration) or TMA_4_PP_i_ (at high substrate concentration) and the P_i_ accumulation was continuously recorded for 2–3 min at 40 °C. Reaction rates were calculated from the initial slopes of the P_i_ liberation and analysed using Prism 6.0 (GraphPad Software) based on a model assuming allosteric substrate binding in dimeric enzyme (Scheme 1 and Equation 1)(Anashkin et al., 2021) or a standard Michaelis-Menten type mechanism.

**Scheme 1:**
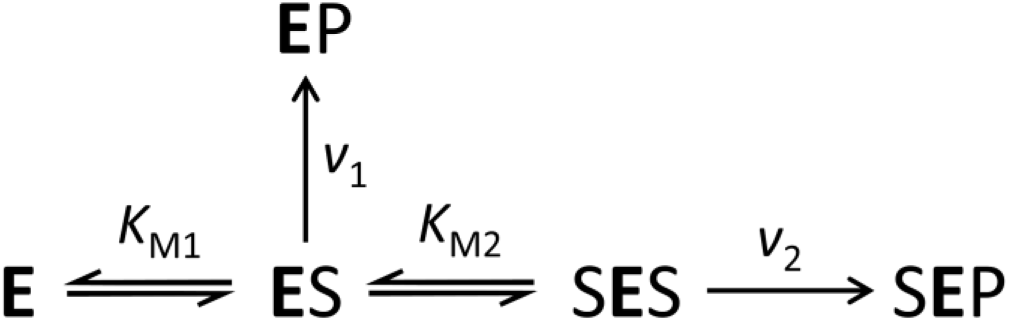
Kinetic model for the catalysis by M-PPase homodimeic enzyme

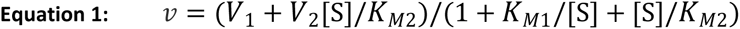

### Nanion SURFE2R measurement

For the Nanion SURFE2R experiments, purified *Tm*PPase was reconstituted into liposomes as previously described (Li et al., 2016) with some modifications. Briefly, the purified protein was buffer exchanged into a reconstitution buffer (50mM MOPS-KOH pH 7.2, 50 mM KCl, 5 mM MgCl_2_, 2 mM DTT) to remove Na^+^ and glycerol and then diluted to 50 μg mL^-1^. 120 mg of soybean lectin was dissolved in 1 mL of water and tip sonicated with 60% amplitude, 6 second pulses for 1 minute with 1 minute on ice between sonications until the solution was clear. 15 μL liposomes solution (120 mg mL^-1^ soybean lecithin in 50 mM MOPS-KOH pH 7.2) was mixed with 1 mL of diluted protein sample. SM-2 Bio-beads were added in small increments to a final concentration of 0.25 mg μL^-1^ and then placed into a mixer at 4 °C for 6 hours to ensure that the beads stayed in suspension. The proteoliposomes were collected and frozen at -80 °C in aliquots. To ensure that the reconstituted protein was still active, the hydrolytic activity was assessed in fixed-time P_i_-release assays as described above.

Electrometric measurements were performed on the SURFE2R N1 instrument (Nanion Technology). The gold sensors were prepared based on the ‘SURFE2R N1 protocol’. This involves thiolating the gold sensor surface and covering it with a lipid layer using sensor prep A2 and B solutions. The resulting solid support membrane-based biosensor can be used to immobilize liposomes containing *Tm*PPase. 15 μl of sonicated proteoliposomes followed by 50ul of *Tm*PPase SURFE2R buffer (50 mM MOPS-KOH pH 7.2, 50 mM NaCl, 5 mM MgCl_2_) were applied directly to the sensor surface. Sensors were centrifuged for 30 mins at 2500 g and incubated at 4 °C for 3 hours. After mounting the sensors in the SURFE2R N1, the sensors were rinsed twice with 1 mL rinsing buffer (50 mM MOPS-KOH pH 7.2, 50 mM NaCl, 5 mM MgCl2). Measurements were performed for 3 seconds by consecutively flowing non-activating buffer B (50 mM MOPS-KOH 7.2, 5 mM MgCl_2_, 200 mM K_2_HPO_4_) and activating buffer A (50 mM MOPS-KOH, 50 mM NaCl, 5 mM MgCl_2_, K_4_PP_i_ or IDP) across the sensor for 1 second each in a BAB sequence. Thus, charge transport across the membrane is initiated by K_4_PP_i_ or IDP in buffer A, which is flowed across the sensor during the time period between 1 to 2 seconds. Transport of positively charged ions during this time results in a positive electrical current, which is the signal output of the SURFE2R N1 instrument.

## Figure supplements

**Figure 2-figure supplement 1:**
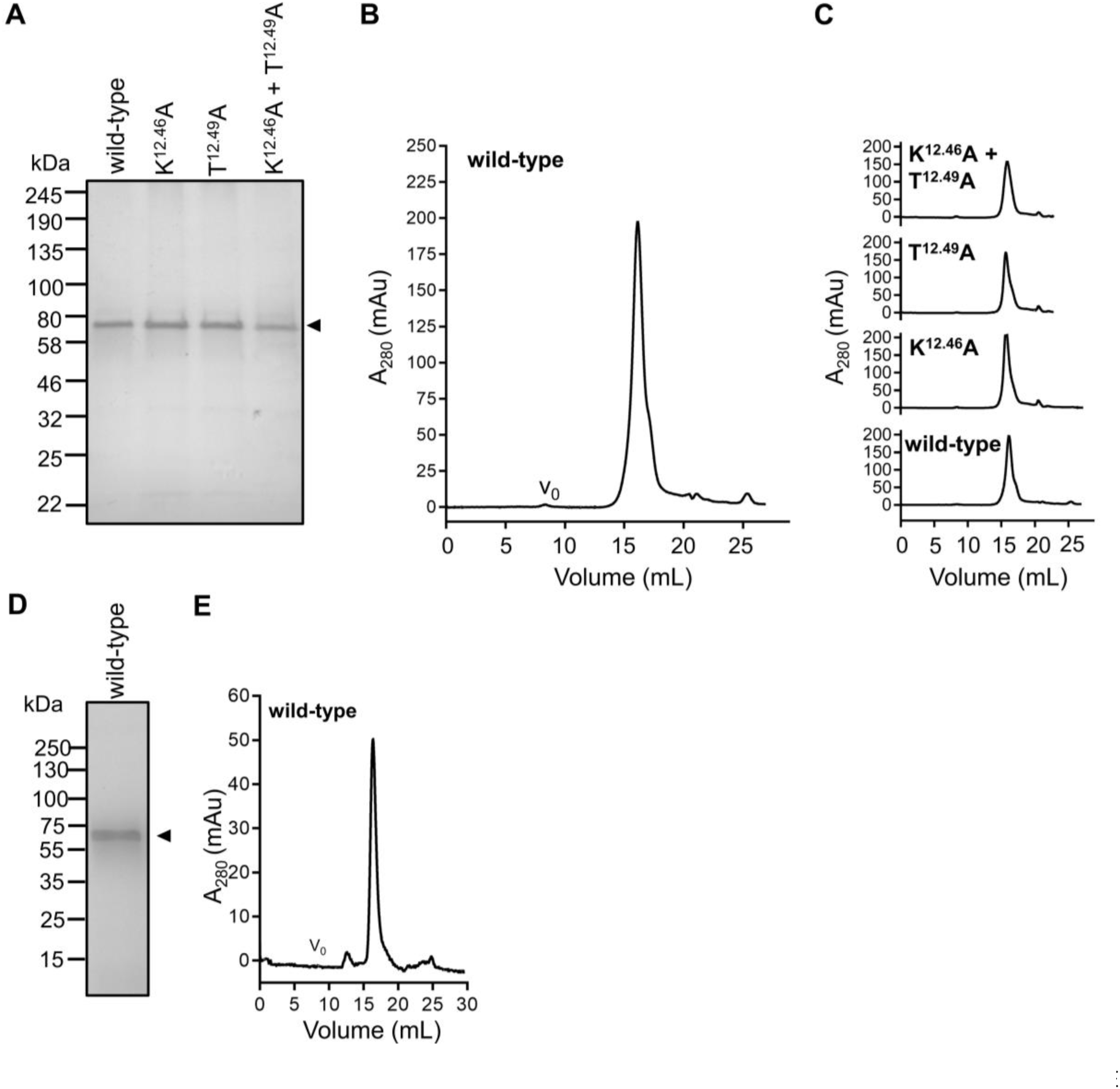
Hot-solve” purification of and analytical SEC of M-PPases. (**A**) SDS-PAGE (Coomassie stain) analysis of purified wild-type and variant *Pa*PPase. (**B**) Analytical SEC of wild-type *Pa*PPase on Superose 6 Increase 10/300 column. The void volume is indicated by vo. (**C**) SEC elution volume comparison of wild-type and variant *Pa*PPase on Superose 6 Increase 10/300 column. (**D**) SDS-PAGE (Coomassie stain) analysis of purified wild-type *Tm*PPase. (**E**) Analytical SEC of wild-type *Tm*PPase on Superose 6 Increase 10/300 column. The void volume is indicated by vo.

**Figure 2-figure supplement 2:**
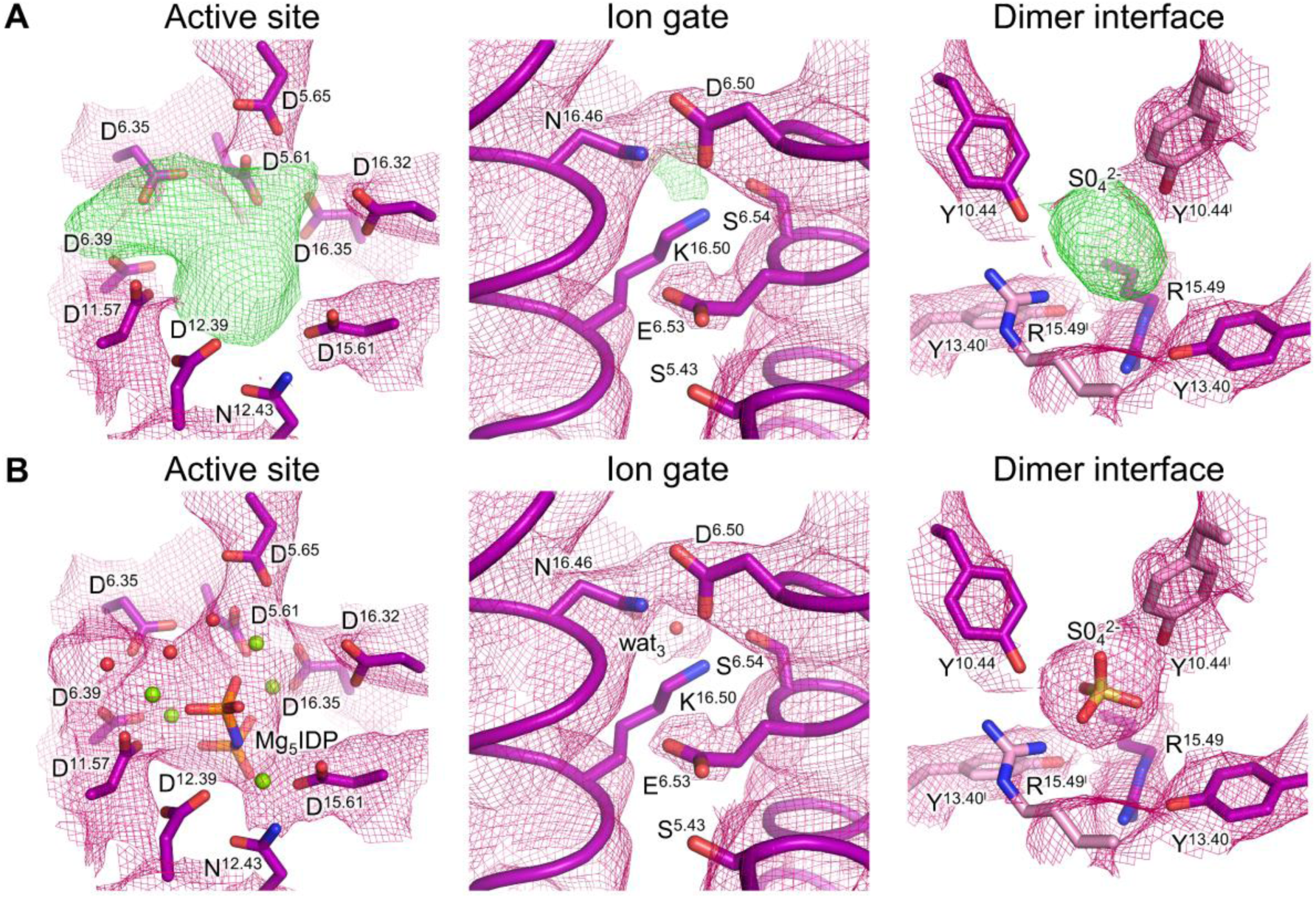
Electron density maps of *Pa*PPase:Mg5IDP key regions. (**A**) 2mFo-dFc map of active site, ion gate and dimer interface residues and mFo-dFc omit map with positive density shown in green and red for ligand or heteroatom binding regions. (**B**) 2mFo-dFc map of active site, ion gate and dimer interface residues with ligands and heteroatoms added to the model. Mg^2+^ are shown as green spheres and structural water molecules are shown as red spheres. Residues of subunit A are coloured in purple and residues of subunit B a coloured in pink (additionally marked with apostrophes).

**Figure 2-figure supplement 3:**
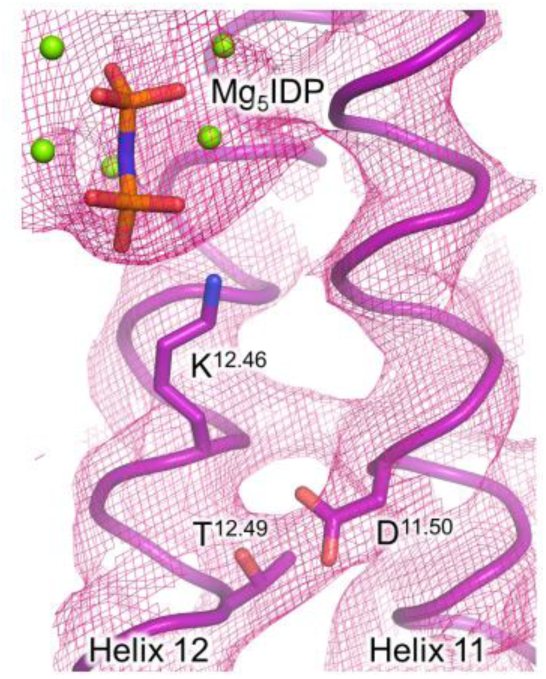
2mFo-dFc: electron density map of the K^+^/K^12.46^ cationic centre of *Pa*PPase:Mg5IDP. 2mFo-dFc: electron density of key residue K^12.46^ and nearby residues T^12.49^ and D^11.50^ are shown at 3 σ.

**Figure 3-figure supplement 1:**
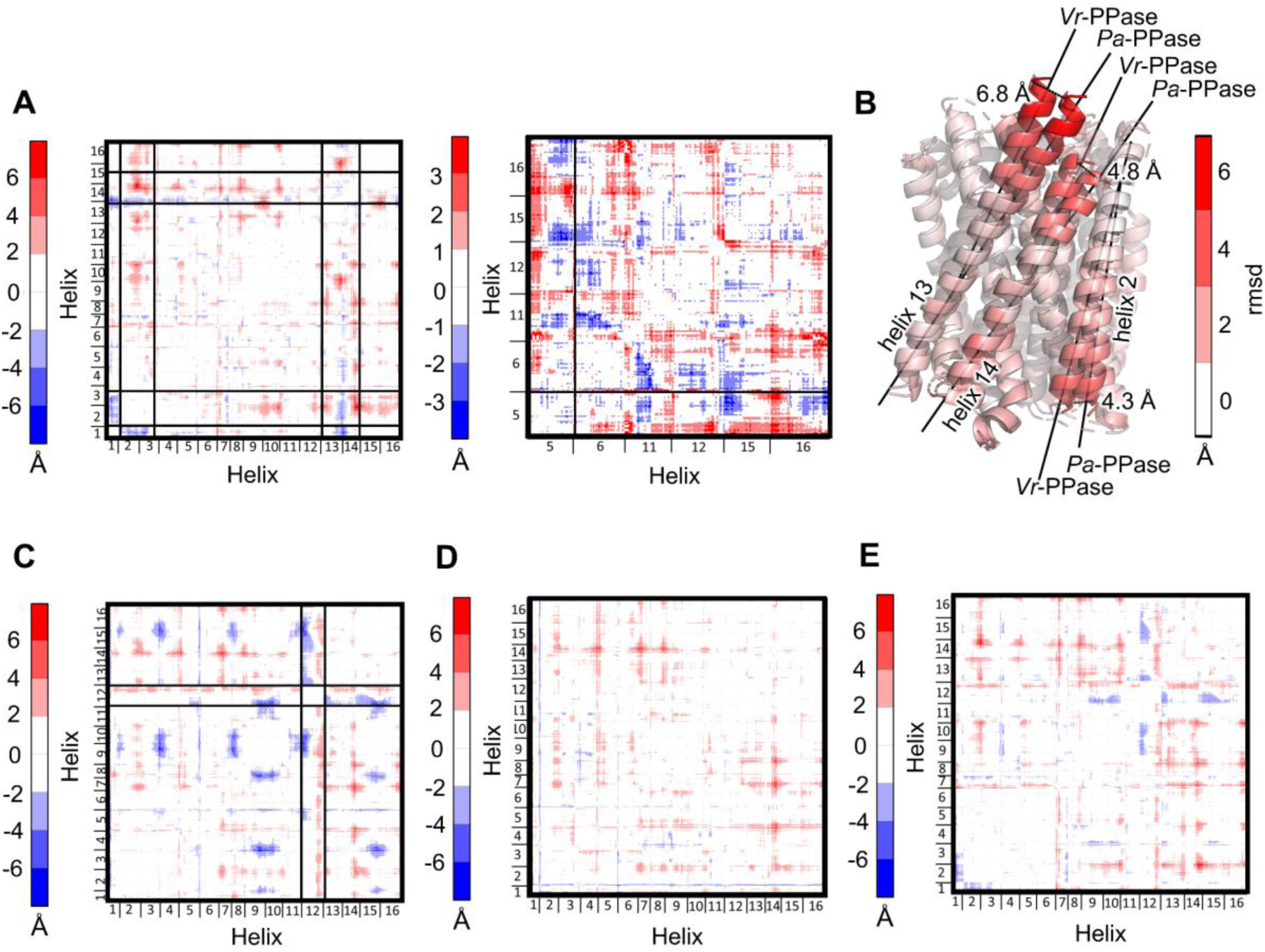
**Comparison of inter-Cα distances between *Pa*PPase:Mg5IDP and other M-PPase structures**. The difference in inter-Cα distances is coloured from red (biggest difference) to blue (smallest difference) in each selection and helices with large clusters of changes are highlighted by black boxes. (**A**) Difference distance matrix (DiDiMa) of *Pa*PPase:Mg5IDP versus *Vr*PPase:Mg5IDP. Left panel shows the DiDiMa of all atoms (scale: ±6 Å); right panel shows inter-atom differences of inner ring helices only (scale: ±3 Å). (**B**) Structural alignment of subunit A of the *Pa* and *Vr*PPase Mg5IDP complexes, with helices coloured by their r.m.s.d./Cα. Dashed lines indicate the distances measured at the end of the helices. (**C-E**) DiDiMa (scale: ±6 Å) of *Pa*PPase:Mg5IDP *versus* (**C**) *Tm*PPase:CaMg; (**D**) *Tm*PPase:Mg4Pi2 and *Vr*PPase:Mg2Pi (**E**).

**Figure 3-figure supplement 2:**
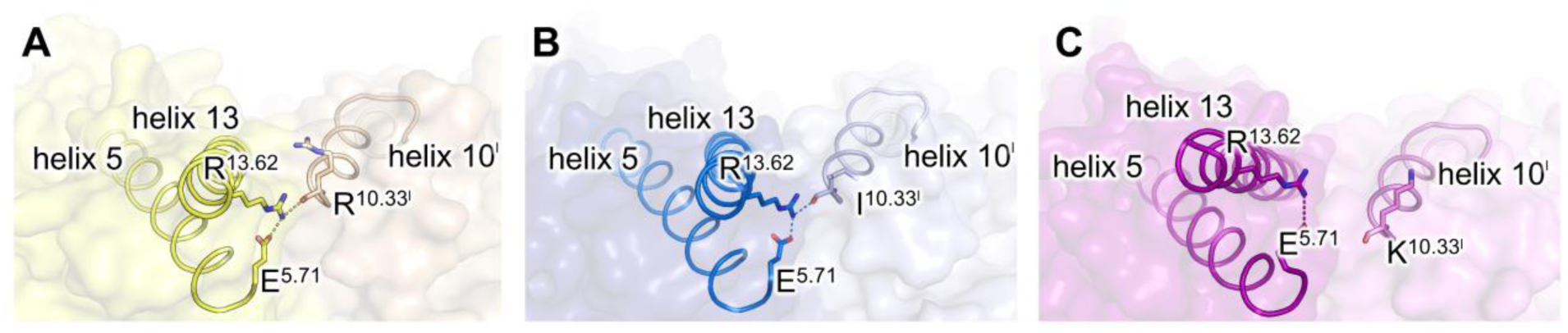
Orientation of helix 5, helix 13 and helix 10 (opposing subunit) in different IDP-bound M-PPase structures. (**A**) Orientation shown in *Vr*PPase:Mg5IDP. (**B**) Orientation shown in *Tm*PPase:Mg5IDP. (**C**) Orientation shown in *Pa*PPase:Mg5IDP. Salt bridge network interactions of E^5.71^-R^13.62^- R/K^10.33’^ are represented by dashed lines in both panels. Subunit A and B are coloured differently, and residues of subunit B are marked by an apostrophe.

**Figure 5-figure supplement 1:**
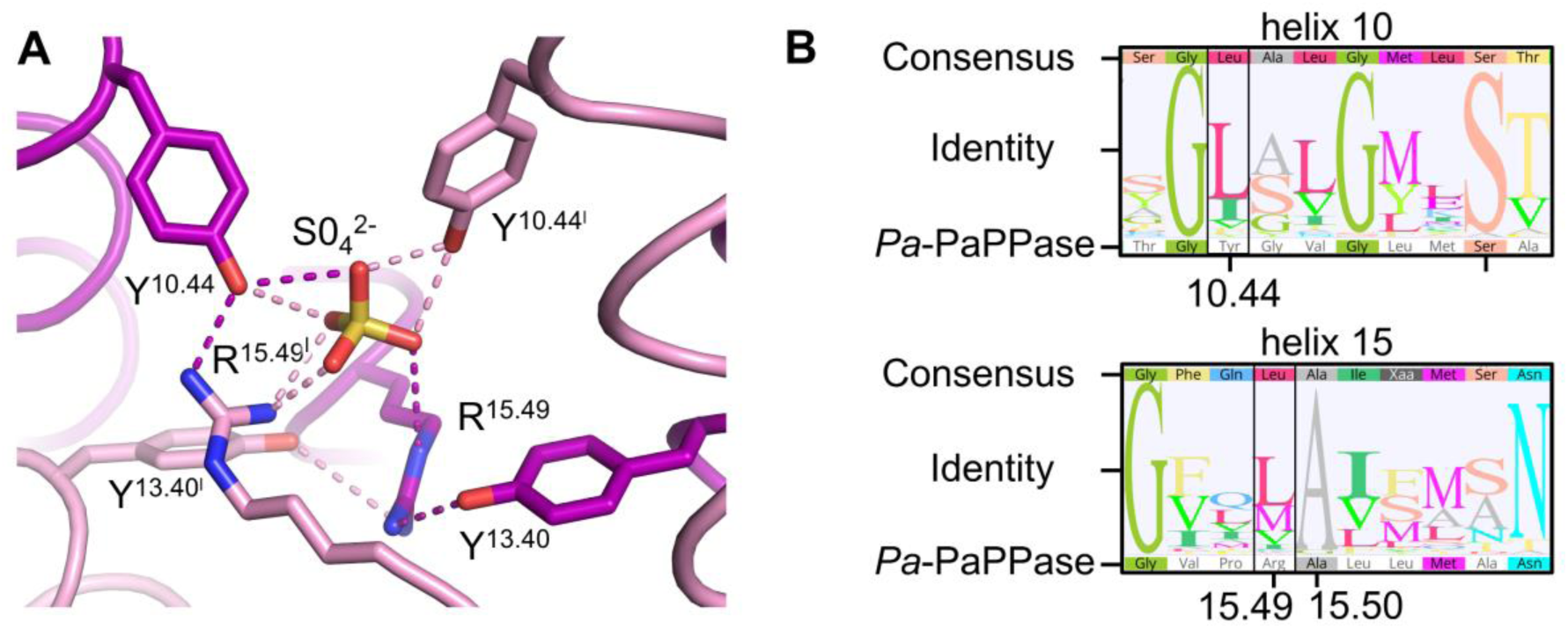
SO4^2-^ binding site at the dimer interface of *Pa*PPase:Mg5IDP. (**A**) Structural overview with subunit A in purple and subunit B in pink (additionally marked with apostrophes). Side chain interactions are shown as dashed lines. (**B**) Sequence analysis of the SO4^2-^ binding site. The consensus sequence and sequence identity (sequence logo showing the graphical representation of the residue conservation) are based on an alignment of 45 homologous sequences to *Pa*PPase identified in a blastp search of the UniProt database. Residues of interest are highlighted by a black box and labelled following the B&W convention.

**Figure 6-figure supplement 1:**
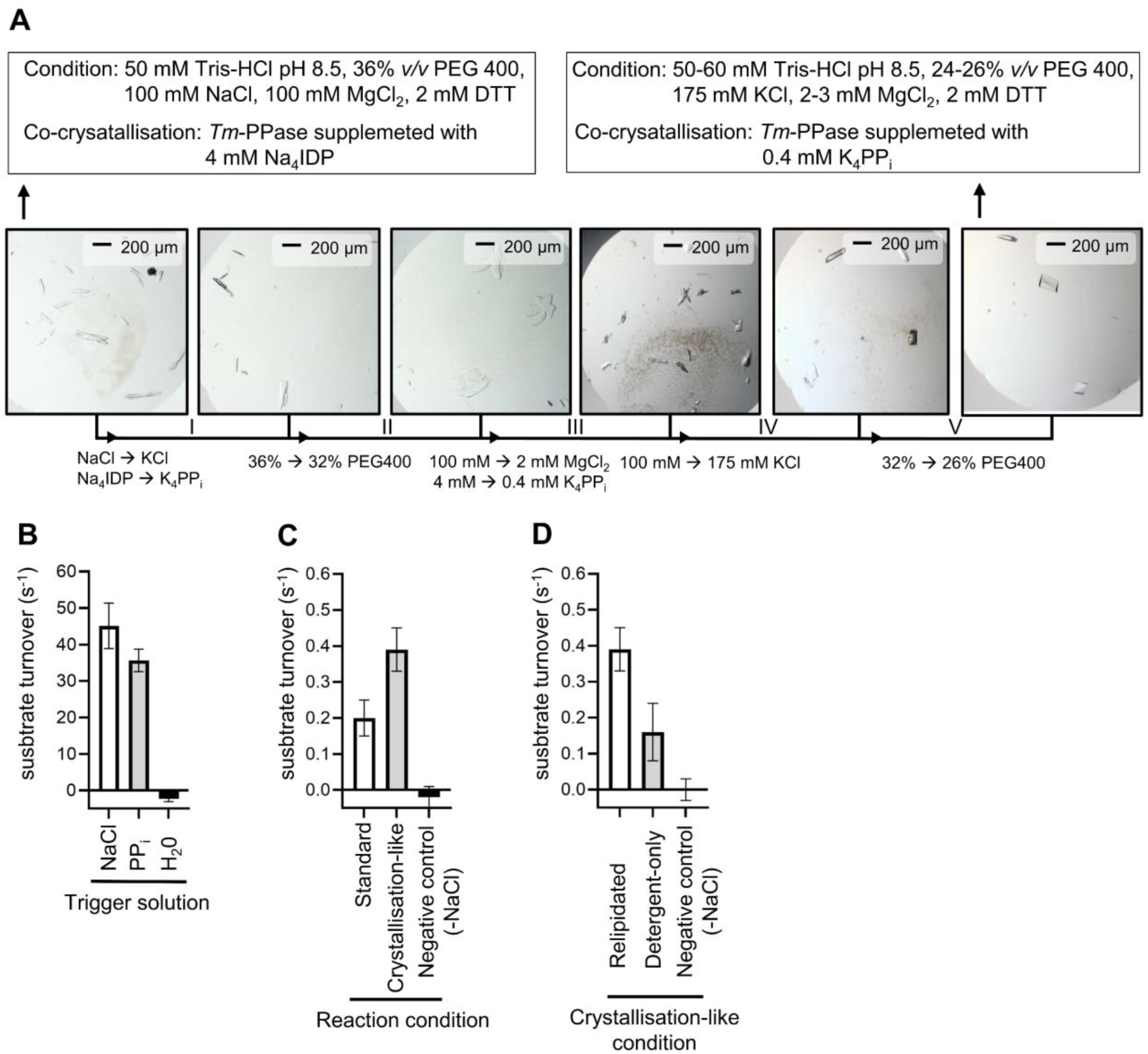
Pre-studies of time-resolved crystallographic experiments with *Tm*PPase. (**A**) Crystal optimisation rounds I-V. Arrows indicate the optimisation steps and changes are annotated. . The start and end condition are displayed in black boxes. (**B-D**) Quantitative Pi-release activity assays of *Tm*PPase in a range of different reaction conditions. (**B**) Substrate turnover by *Tm*PPase at 71 °C upon reaction initiation with different trigger solutions. (**C**) Substrate turnover by *Tm*PPase at 20 °C in different reaction conditions. (**D**) Substrate turnover by *Tm*PPase at 20 °C in crystallisation-like reaction conditions after different sample treatments. Negative controls lack NaCl in the final reaction condition (-NaCl).

**Figure 6-figure supplement 2:**
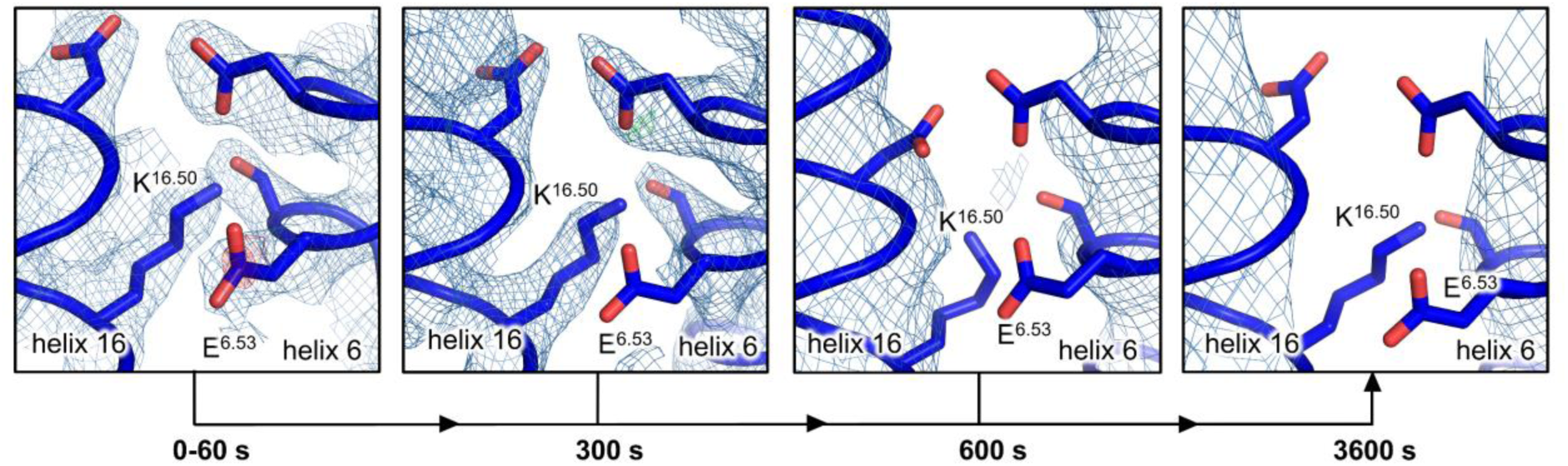
Ion gate of time-resolved *Tm*PPase structures. Structures shown for grouped datasets of different time-points (0-60 seconds) and combined datasets of the same time-point (300, 600 and 3600 seconds). Structures are shown with 2mFo-dFc density (blue) and mFo-dFc density (red/green) at 1 σ and 3 σ, respectively.

**Figure 9-figure supplement 1:**
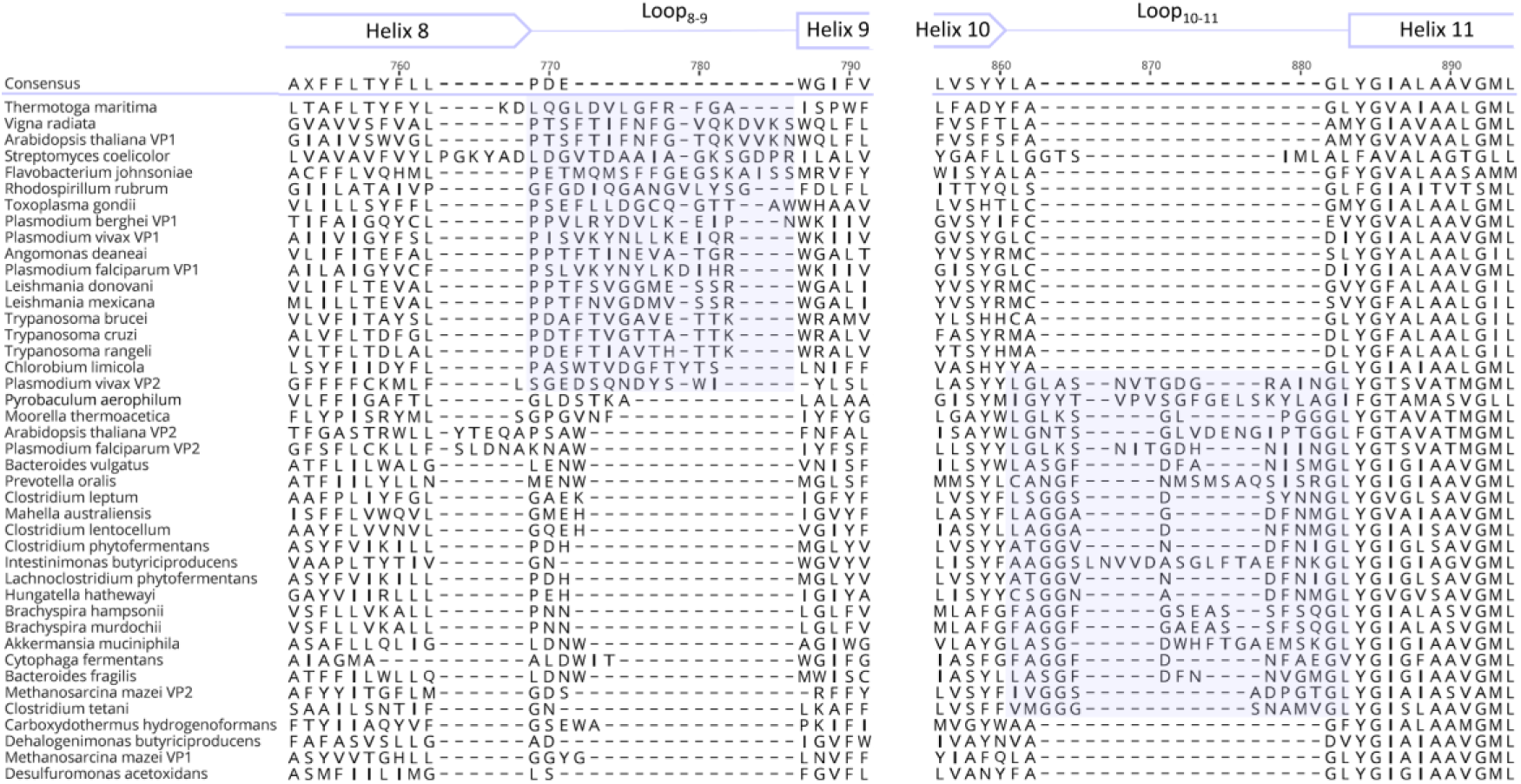
Sequence alignment of exit channel loop regions of a blastp search obtained with *Pa*PPase as search sequence. Sequences are sorted in two blocks (blue box) based on their loop8-9 and loop10-11 length. Annotation of topology is based on the *Tm*PPase:CaMg structure and may differ in non-conserved regions.

## Additional files

**Supplementary table 1:** M-PPase numbering scheme.

| B&W<sup>‡</sup> | *Tm*PPase | *Vr*PPase | *Pa*PPase |
| --- | --- | --- | --- |
| 1.50 | F17 | F25 | Y20 |
| 2.50 | K55 | K94 | R58 |
| 3.50 | S87 | S153 | S96 |
| 4.50 | G130 | G194 | G138 |
| 5.43 | S184 | S235 | S176 |
| 5.50 | R191 | R242 | R183 |
| 6.50 | D243 | D294 | D235 |
| 6.53 | E246 | G297 | E238 |
| 6.57 | G250 | R301 | V242 |
| 7.50 | G294 | G335 | A271 |
| 8.50 | L321 | L361 | I294 |
| 9.50 | G369 | G411 | G333 |
| 10.50 | S416 | S458 | S380 |
| 11.50 | D458 | D500 | D439 |
| 12.46 | A495 | A537 | K476 |
| 12.49 | G498 | G540 | T479 |
| 12.50 | K499 | K541 | K480 |
| 13.50 | V566 | V597 | V554 |
| 14.50 | M611 | M642 | F599 |
| 15.50 | A649 | A680 | A637 |
| 16.50 | K707 | K742 | K691 |

‡ Ballesteros & Weinstein nomenclature (Ballesteros & Weinstein, 1995)

**Supplementary table 2:** X-ray data collection and refinement statistics of 3.8 Å *Pa*PPase structure.

| Data collection |  |
| --- | --- |
| Space group | P2<sub>1</sub> |
| Cell dimensions |  |
| *a*, *b*, *c* (Å) | 107.2, 88.0, 116.8 |
| $\alpha$, $\beta$, $\gamma$ (°) | 90.0, 106.9, 90.0 |
| Source | DLS: i04/i24 |
|  | ESRF: ID23-1/MASSIF1 |
| Wavelength* (Å) | 0.966/0.979/0.971/0.968 |
| Resolution (Å) | 19.97-3.84 (3.98-3.84) |
| Overall (Å) | 3.8 |
| along *h* axis | 5.3 |
| along *k* axis | 4.1 |
| along *l* axis | 3.8 |
| Measured reflections | 65003 (2063) |
| Unique reflections | 13069 (653) |
| Completeness (%) | 87.6 |
| CC<sub>1/2</sub> | 0.979 |
| Mean *I*/σ(*I*) | 3.9 |
| Multiplicity | 5.0 |
| B-factors (Å<sup>2</sup>) | 112.11 |
| R<sub>merge</sub> | 0.335 (1.524) |
| R<sub>meas</sub> | 0.375 (1.749) |
| R<sub>pim</sub> | 0.163 (0.827) |
| Refinement |  |
| R<sub>work</sub>(%)/R<sub>free</sub>(%) | 28.9/31.1 |
| No. of atoms | 9734 |
| Protein | 9694 |
| Ligands | 33 |
| Water | 7 |
| No. of chains (ASU) | 2 |
| B-factors (Å<sup>2</sup>) | 106.96 |
| Protein | 106.98 |
| Ligands/Ions | 94.85F |
| R. M. S. Deviations |  |
| Bond lengths (Å) | 0.002 |
| Bond angle (°) | 0.047 |
| Ramachandran statistics (%) |  |
| Favoured | 96.61 |
| Allowed | 3.16 |
| Outliers | 0.23 |
| Statistics for the highest-resolution shell are shown in parentheses |  |
| * Data from several beamlines |  |

**Supplementary table 3:**
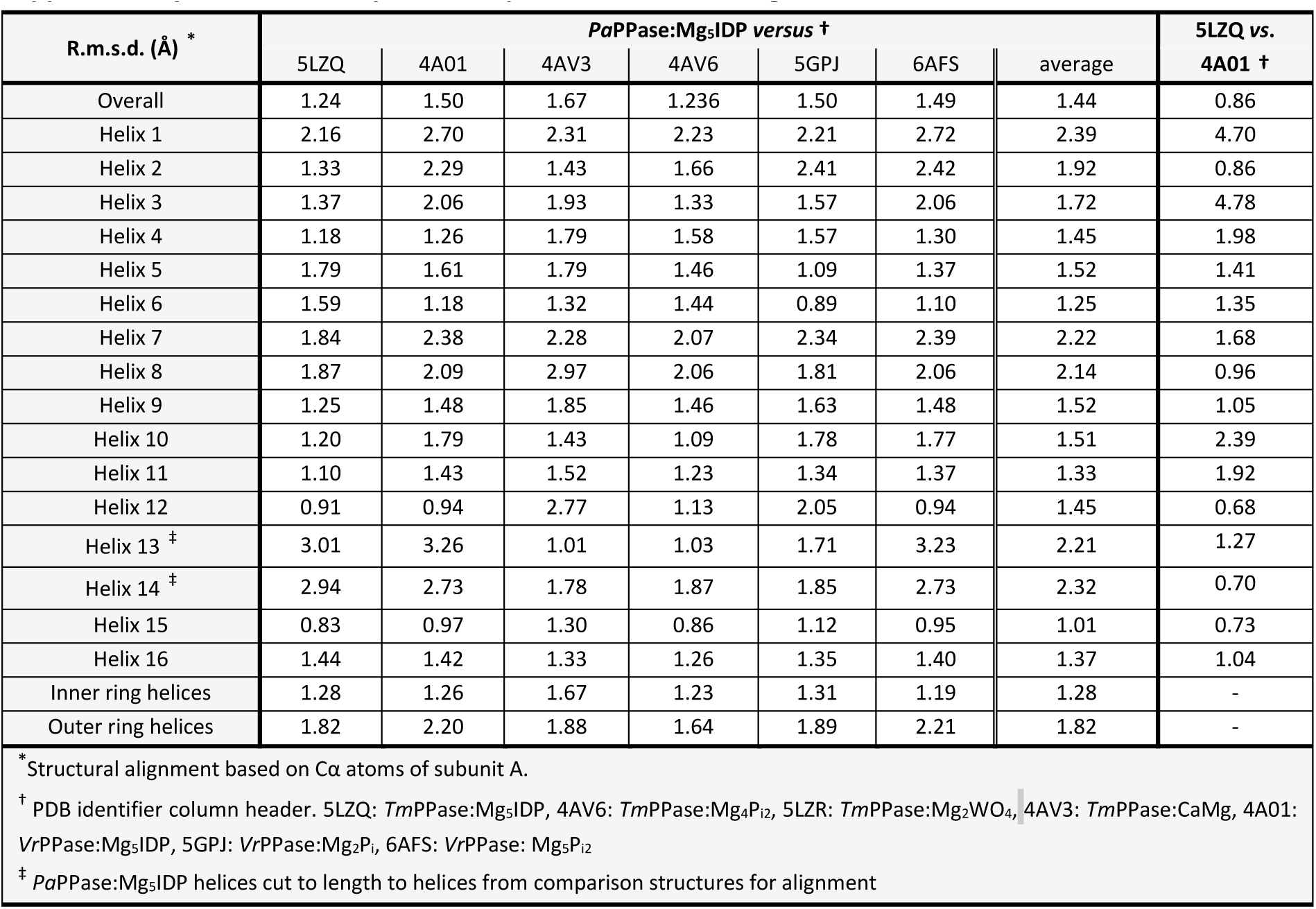
Helix by helix comparison of *Pa*PPase:Mg5IDP structure to other M-PPase structures.

| R.m.s.d. (Å) * | *Pa*PPase:Mg<sub>5</sub>IDP versus † |  |  |  |  |  |  | 5LZQ vs. 4A01 † |
| --- | --- | --- | --- | --- | --- | --- | --- | --- |
|  | 5LZQ | 4A01 | 4AV3 | 4AV6 | 5GPJ | 6AFS | average |  |
| Overall | 1.24 | 1.50 | 1.67 | 1.236 | 1.50 | 1.49 | 1.44 | 0.86 |
| Helix 1 | 2.16 | 2.70 | 2.31 | 2.23 | 2.21 | 2.72 | 2.39 | 4.70 |
| Helix 2 | 1.33 | 2.29 | 1.43 | 1.66 | 2.41 | 2.42 | 1.92 | 0.86 |
| Helix 3 | 1.37 | 2.06 | 1.93 | 1.33 | 1.57 | 2.06 | 1.72 | 4.78 |
| Helix 4 | 1.18 | 1.26 | 1.79 | 1.58 | 1.57 | 1.30 | 1.45 | 1.98 |
| Helix 5 | 1.79 | 1.61 | 1.79 | 1.46 | 1.09 | 1.37 | 1.52 | 1.41 |
| Helix 6 | 1.59 | 1.18 | 1.32 | 1.44 | 0.89 | 1.10 | 1.25 | 1.35 |
| Helix 7 | 1.84 | 2.38 | 2.28 | 2.07 | 2.34 | 2.39 | 2.22 | 1.68 |
| Helix 8 | 1.87 | 2.09 | 2.97 | 2.06 | 1.81 | 2.06 | 2.14 | 0.96 |
| Helix 9 | 1.25 | 1.48 | 1.85 | 1.46 | 1.63 | 1.48 | 1.52 | 1.05 |
| Helix 10 | 1.20 | 1.79 | 1.43 | 1.09 | 1.78 | 1.77 | 1.51 | 2.39 |
| Helix 11 | 1.10 | 1.43 | 1.52 | 1.23 | 1.34 | 1.37 | 1.33 | 1.92 |
| Helix 12 | 0.91 | 0.94 | 2.77 | 1.13 | 2.05 | 0.94 | 1.45 | 0.68 |
| Helix 13 ‡ | 3.01 | 3.26 | 1.01 | 1.03 | 1.71 | 3.23 | 2.21 | 1.27 |
| Helix 14 ‡ | 2.94 | 2.73 | 1.78 | 1.87 | 1.85 | 2.73 | 2.32 | 0.70 |
| Helix 15 | 0.83 | 0.97 | 1.30 | 0.86 | 1.12 | 0.95 | 1.01 | 0.73 |
| Helix 16 | 1.44 | 1.42 | 1.33 | 1.26 | 1.35 | 1.40 | 1.37 | 1.04 |
| Inner ring helices | 1.28 | 1.26 | 1.67 | 1.23 | 1.31 | 1.19 | 1.28 | - |
| Outer ring helices | 1.82 | 2.20 | 1.88 | 1.64 | 1.89 | 2.21 | 1.82 | - |

\* Structural alignment based on Cα atoms of subunit A.
† PDB identifier column header. 5LZQ: *Tm*PPase:Mg<sub>5</sub>IDP, 4AV6: *Tm*PPase:Mg<sub>4</sub>Pi<sub>2</sub>, 5LZR: *Tm*PPase:Mg<sub>2</sub>WO<sub>4</sub>, 4AV3: *Tm*PPase:CaMg, 4A01: *Vr*PPase:Mg<sub>5</sub>IDP, 5GPJ: *Vr*PPase:Mg<sub>2</sub>Pi, 6AFS: *Vr*PPase: Mg<sub>5</sub>Pi<sub>2</sub>
‡ *Pa*PPase:Mg<sub>5</sub>IDP helices cut to length to helices from comparison structures for alignment

**Supplementary table 4:** HELANAL-Plus curvature analysis of helix 5 of *Pa*PPase:Mg5IDP.

| HELANAL-Plus Parameters * | Helix 5<br>*Pa*PPase:Mg<sub>5</sub>IDP | Helix 5<br>*Vr*PPase:Mg<sub>5</sub>IDP |
| --- | --- | --- |
| Helix length (residues) | 37 | 37 |
| Average number of residues per turn | 3.66 | 3.69 |
| Average unit height of helix (Å) | 1.5 | 1.53 |
| Average virtual torsion angle (°) | 49.8 | 49.8 |
| Average bending angle (°) | 11.9 | 11 |
| Maximum bending angle (°) | 24.5 | 23.2 |
| Radius of sphere curvature (Å) | 114 | 76 |
| R.m.s.d. of sphere fit (Å) (r.m.s.d.S) | 0.247 | 0.266 |
| R.m.s.d. of linear fit (Å) (r.m.s.d.L) | 0.154 | 0.238 |
| Geometry† | linear | kinked |

\* Definition of all parameters can be found in (Kumar & Bansal, 2012).
† Classification: Linear if, r.m.s.d.S > r.m.s.d.L and maximum bending angle (MBA) < 20° and if the 20° < MBA < 30°, r.m.s.d.S and r.m.s.d.L are both < 0.14 and 0.16 Å, respectively, and r.m.s.d.S > r.m.s.d.L. Kinked, if the value of the MBA is > 30° or the MBA is between 20° and 30° and r.m.s.d. to sphere (r.m.s.d.S) and 3D line (r.m.s.d.L) fit is more than 0.14 and 0.16 Å, respectively.

**Supplementary table 5:** Comparison of the hydrogen bonding pattern around S^5.43^ and D^6.43^ in IDP-bound structures.

| Residue from | Helical geometry* |  |  |
| --- | --- | --- | --- |
|  | *Pa*PPase:Mg<sub>5</sub>IDP | *Vr*PPase:Mg<sub>5</sub>IDP | *Tm*PPase:Mg<sub>5</sub>IDP |
| 5.37 | (α) | 3<sub>10</sub> | 3<sub>10</sub> |
| 5.38 | 3<sub>10</sub> | α | 3<sub>10</sub> |
| 5.39 | 3<sub>10</sub> | α | (α) |
| 5.40 | (α) | α | (α) |
| 5.41 | α | α | α |
| 5.42 | 3<sub>10</sub> | α | α |
| 5.43 | α | α | α |
| 5.43 | 3<sub>10</sub> | 3<sub>10</sub> | α |
| 5.44 | α | α | α |
| 5.45 | (α) | α | α |
| 5.46 | α | α | α |
| 6.40 | α | (α) | 3<sub>10</sub> |
| 6.41 | 3<sub>10</sub> | α | 3<sub>10</sub> |
| 6.42 | α | α | α |
| 6.43 | α | (α) | (α) |
| 6.44 | (α) | π | π |
| 6.45 | π | π | π |
| 6.46 | π | α | α |
| 6.47 | α | 3<sub>10</sub> | π |
| * Hydrogen bonding from amino nitrogen to upstream carbonyl oxygen of main chain |  |  |  |
| Entries in parentheses are based on rise per residue as no hydrogen bond (distance > 4 Å) is formed |  |  |  |

**Supplementary table 6:** Rotamer options for K^12.46^ in *Pa*PPase:Mg5IDP of the backbone- independent Richardson library. Sorting based on their vdW radii overlap to surrounding atoms from lowest to highest. Modelled rotamer at the top of the table.

| Chi<sub>1</sub> | Chi<sub>2</sub> | Chi<sub>3</sub> | Chi<sub>4</sub> | Sum vdW radii overlap [Å] | Clashes* | Hydrogen bonds |
| --- | --- | --- | --- | --- | --- | --- |
| 62 | 180 | 68 | 180 | 0.62 | 1 | 3 |
| 62 | 180 | -68 | 180 | 2.00 | 2 | 4 |
| 63 | -178 | 178 | -179 | 2.55 | 3 | 2 |
| 63 | -170 | -177 | 72 | 2.84 | 3 | 1 |
| 62 | 180 | 180 | -65 | 3.03 | 3 | 1 |
| -70 | -179 | -66 | -64 | 4.59 | 4 | 2 |
| -70 | -170 | -66 | -175 | 5.96 | 5 | 3 |
| 179 | 59 | 163 | 60 | 7.35 | 7 | 0 |
| -177 | 180 | 68 | 65 | 7.46 | 7 | 1 |
| -59 | -69 | -176 | -70 | 7.53 | 8 | 3 |
| -58 | -61 | -177 | -179 | 8.31 | 10 | 2 |
| -69 | 164 | 62 | -179 | 8.44 | 7 | 2 |
| 180 | 179 | 78 | 179 | 8.50 | 9 | 4 |
| -67 | -176 | 174 | 76 | 9.04 | 8 | 0 |
| -177 | 178 | 179 | 180 | 9.35 | 9 | 1 |
| -177 | 68 | 180 | -65 | 9.62 | 8 | 1 |
| -62 | -68 | 180 | 65 | 9.74 | 10 | 1 |
| -177 | 180 | 171 | 63 | 10.01 | 10 | 2 |
| -90 | 68 | 180 | 180 | 10.23 | 10 | 2 |
| 179 | 172 | 178 | -72 | 10.36 | 8 | 1 |
| -177 | 62 | 173 | 171 | 11.62 | 11 | 2 |
| -67 | -179 | -179 | -63 | 12.45 | 11 | 1 |
| -59 | -58 | -75 | -174 | 12.62 | 12 | 1 |
| -175 | -174 | -69 | 179 | 12.94 | 10 | 2 |
| The chi angles translate to m for minus (-60°), t for trans (±180°) and p for plus (+60°) per chi angle in Coot |  |  |  |  |  |  |
| * vdW radii overlap of ≥ 0.6 Å is classified as a clash |  |  |  |  |  |  |

**Supplementary table 7:** Rotamer options for K^16.50^ *Pa*PPase:Mg5IDP of the backbone-independent Richardson library. Sorting based on their vdW radii overlap to surrounding atoms from lowest to highest. Modelled rotamer at the top of the table.

| Chi<sub>1</sub> | Chi<sub>2</sub> | Chi<sub>3</sub> | Chi<sub>4</sub> | Sum vdW radii overlap [Å] | Clashes* | Hydrogen bonds |
| --- | --- | --- | --- | --- | --- | --- |
| -70 | -170 | -66 | -175 | 0 | 0 | 4 |
| -69 | 164 | 62 | -179 | 0 | 0 | 1 |
| -69 | -179 | 70 | 67 | 0 | 0 | 0 |
| -177 | 180 | -68 | -65 | 0.83 | 1 | 0 |
| -175 | -174 | -69 | 179 | 2.17 | 3 | 0 |
| -70 | -179 | -66 | -64 | 3.08 | 3 | 2 |
| -59 | -58 | -75 | -174 | 3.97 | 3 | 1 |
| -67 | 176 | 179 | 177 | 6.41 | 5 | 2 |
| -177 | 68 | 180 | -65 | 8.98 | 9 | 1 |
| 62 | 180 | 68 | 180 | 9.03 | 7 | 4 |
| -67 | -179 | -179 | -63 | 9.1 | 8 | 0 |
| -67 | -176 | 174 | 76 | 10.38 | 9 | 2 |
| -177 | 178 | 179 | 180 | 10.63 | 11 | 1 |
| -177 | 180 | 68 | 65 | 11.13 | 7 | 1 |
| 179 | 62 | 173 | 171 | 11.68 | 12 | 0 |
| -62 | -68 | 180 | 65 | 12.41 | 10 | 1 |
| 179 | 59 | 163 | 60 | 12.62 | 12 | 0 |
| -177 | 172 | 178 | -72 | 12.77 | 11 | 1 |
| -177 | 180 | 171 | 63 | 12.85 | 12 | 0 |
| -58 | -61 | -177 | -179 | 13.73 | 10 | 2 |
| -59 | -69 | -176 | -70 | 15.19 | 11 | 0 |
| 180 | 179 | 78 | 179 | 15.78 | 11 | 0 |
| 63 | -178 | 178 | -179 | 17.19 | 14 | 1 |
| 62 | 180 | 180 | -65 | 19.7 | 15 | 1 |
| The chi angles translate to m for minus (-60°), t for trans (±180°) and p for plus (+60°) per chi angle in Coot |  |  |  |  |  |  |
| * vdW radii overlap of ≥ 0.6 Å is classified as a clash |  |  |  |  |  |  |

**Supplementary table 8:** Catalytic turnover of wild-type *Tm*PPase used for time-resolved structural studies in various conditions upon NaCl activation.

| Fixed-time P<sub>i</sub> release assays | Wild-type TmPPase |  |  |  |
| --- | --- | --- | --- | --- |
| Reaction condition | Standard | Standard | Crystallisation-like | Crystallisation-like |
| Sample treatment | Relipidated | Relipidated | Relipidated | Detergent-only |
| Temperature (°C) | 71 | 20 | 20 | 20 |
| k<sub>cat</sub> (s<sup>-1</sup>)* | 45.13±3.59 | 0.20±0.03 | 0.39±0.03 | 0.16±0.05 |
| * Error is shown as SEM of three technical repeats. |  |  |  |  |

**Supplementary table 9:**
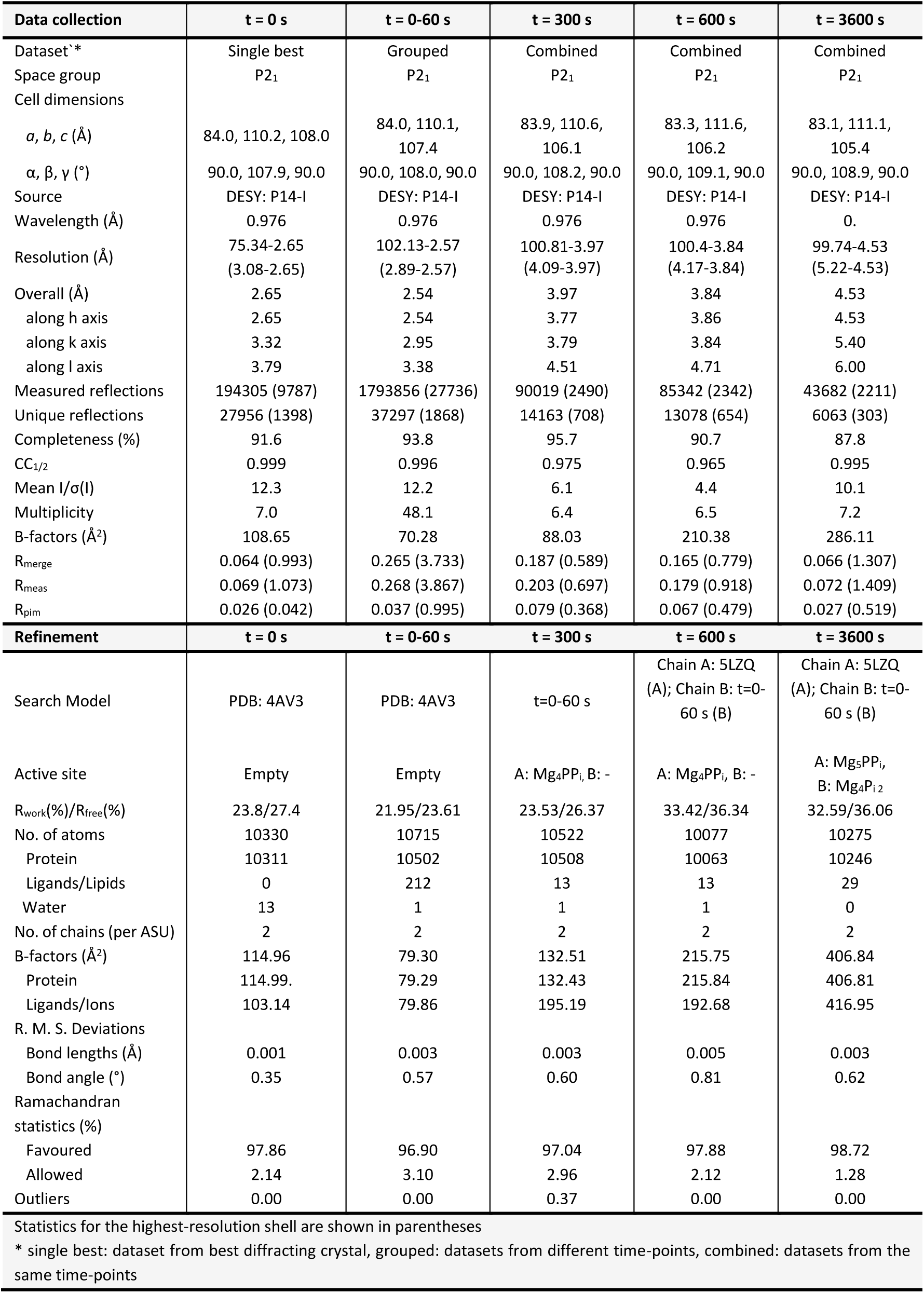
X-ray data collection and refinement statistics of time-resolved *Tm*PPase structures at 0, 0-60, 300, 600 and 3600 seconds post-activation.

| Data collection | t = 0 s | t = 0-60 s | t = 300 s | t = 600 s | t = 3600 s |
| --- | --- | --- | --- | --- | --- |
| Dataset* | Single best | Grouped | Combined | Combined | Combined |
| Space group | P2<sub>1</sub> | P2<sub>1</sub> | P2<sub>1</sub> | P2<sub>1</sub> | P2<sub>1</sub> |
| Cell dimensions |  |  |  |  |  |
| a, b, c (Å) | 84.0, 110.2, 108.0 | 84.0, 110.1,<br>107.4 | 83.9, 110.6,<br>106.1 | 83.3, 111.6,<br>106.2 | 83.1, 111.1,<br>105.4 |
| α, β, γ (°) | 90.0, 107.9, 90.0 | 90.0, 108.0, 90.0 | 90.0, 108.2, 90.0 | 90.0, 109.1, 90.0 | 90.0, 108.9, 90.0 |
| Source | DESY: P14-I | DESY: P14-I | DESY: P14-I | DESY: P14-I | DESY: P14-I |
| Wavelength (Å) | 0.976 | 0.976 | 0.976 | 0.976 | 0. |
| Resolution (Å) | 75.34-2.65<br>(3.08-2.65) | 102.13-2.57<br>(2.89-2.57) | 100.81-3.97<br>(4.09-3.97) | 100.4-3.84<br>(4.17-3.84) | 99.74-4.53<br>(5.22-4.53) |
| Overall (Å) | 2.65 | 2.54 | 3.97 | 3.84 | 4.53 |
| along h axis | 2.65 | 2.54 | 3.77 | 3.86 | 4.53 |
| along k axis | 3.32 | 2.95 | 3.79 | 3.84 | 5.40 |
| along l axis | 3.79 | 3.38 | 4.51 | 4.71 | 6.00 |
| Measured reflections | 194305 (9787) | 1793856 (27736) | 90019 (2490) | 85342 (2342) | 43682 (2211) |
| Unique reflections | 27956 (1398) | 37297 (1868) | 14163 (708) | 13078 (654) | 6063 (303) |
| Completeness (%) | 91.6 | 93.8 | 95.7 | 90.7 | 87.8 |
| CC<sub>1/2</sub> | 0.999 | 0.996 | 0.975 | 0.965 | 0.995 |
| Mean I/σ(I) | 12.3 | 12.2 | 6.1 | 4.4 | 10.1 |
| Multiplicity | 7.0 | 48.1 | 6.4 | 6.5 | 7.2 |
| B-factors (Å<sup>2</sup>) | 108.65 | 70.28 | 88.03 | 210.38 | 286.11 |
| R<sub>merge</sub> | 0.064 (0.993) | 0.265 (3.733) | 0.187 (0.589) | 0.165 (0.779) | 0.066 (1.307) |
| R<sub>meas</sub> | 0.069 (1.073) | 0.268 (3.867) | 0.203 (0.697) | 0.179 (0.918) | 0.072 (1.409) |
| R<sub>pim</sub> | 0.026 (0.042) | 0.037 (0.995) | 0.079 (0.368) | 0.067 (0.479) | 0.027 (0.519) |
| Refinement | t = 0 s | t = 0-60 s | t = 300 s | t = 600 s | t = 3600 s |
| Search Model | PDB: 4AV3 | PDB: 4AV3 | t=0-60 s | Chain A: 5LZQ<br>(A); Chain B: t=0-60 s (B) | Chain A: 5LZQ<br>(A); Chain B: t=0-60 s (B) |
| Active site | Empty | Empty | A: Mg<sub>4</sub>PP<sub>i</sub>, B: - | A: Mg<sub>4</sub>PP<sub>i</sub>, B: - | A: Mg<sub>5</sub>PP<sub>i</sub>,<br>B: Mg<sub>4</sub>P<sub>i2</sub> |
| R<sub>work</sub>(%)/R<sub>free</sub>(%) | 23.8/27.4 | 21.95/23.61 | 23.53/26.37 | 33.42/36.34 | 32.59/36.06 |
| No. of atoms | 10330 | 10715 | 10522 | 10077 | 10275 |
| Protein | 10311 | 10502 | 10508 | 10063 | 10246 |
| Ligands/Lipids | 0 | 212 | 13 | 13 | 29 |
| Water | 13 | 1 | 1 | 1 | 0 |
| No. of chains (per ASU) | 2 | 2 | 2 | 2 | 2 |
| B-factors (Å<sup>2</sup>) | 114.96 | 79.30 | 132.51 | 215.75 | 406.84 |
| Protein | 114.99 | 79.29 | 132.43 | 215.84 | 406.81 |
| Ligands/Ions | 103.14 | 79.86 | 195.19 | 192.68 | 416.95 |
| R. M. S. Deviations |  |  |  |  |  |
| Bond lengths (Å) | 0.001 | 0.003 | 0.003 | 0.005 | 0.003 |
| Bond angle (°) | 0.35 | 0.57 | 0.60 | 0.81 | 0.62 |
| Ramachandran<br>statistics (%) |  |  |  |  |  |
| Favoured | 97.86 | 96.90 | 97.04 | 97.88 | 98.72 |
| Allowed | 2.14 | 3.10 | 2.96 | 2.12 | 1.28 |
| Outliers | 0.00 | 0.00 | 0.37 | 0.00 | 0.00 |
| Statistics for the highest-resolution shell are shown in parentheses |  |  |  |  |  |
| * single best: dataset from best diffracting crystal, grouped: datasets from different time-points, combined: datasets from the same time-points |  |  |  |  |  |

**Supplementary table 10:**
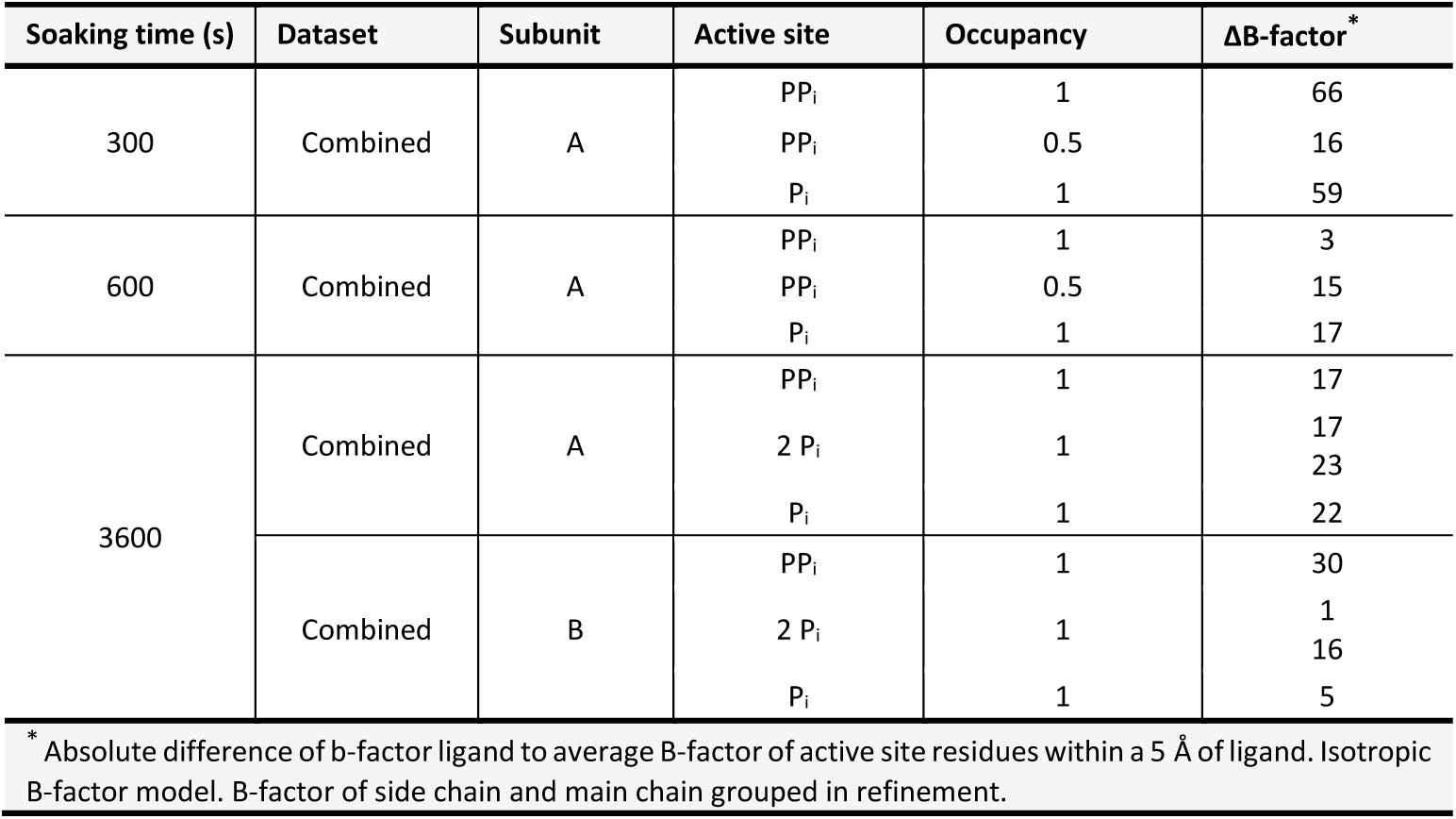
Local B-factor distribution at active site of time-resolved *Tm*PPase structures from different time-points.

## Acknowledgements

We thank the Leeds Astbury Centre for Molecular and Structural Biology for support, Diamond Light Source for access to beam line I04 and I24, the European Synchrotron Radiation Facility for access to beam line ID23-1 and MASSIF-1, and EMBL for access to beam line P14.I and P14.II(T-REXX) at PETRA III (mx747, mx839, mx862).

## Contributions

**Jannik Strauss**

Astbury Centre for Structural and Molecular Biology, University of Leeds, LS2 9JT, Leeds, UK.

Contribution: Conceptualization; Methodology; Validation; Formal analysis; Investigation; Writing – original draft preparation; Writing – review & editing; Visualization; Project administration.

Competing interests: No competing interests declared.

**Craig Wilkinson**

Astbury Centre for Structural and Molecular Biology, University of Leeds, LS2 9JT, Leeds, UK.

Contribution: Conceptualization; Methodology; Validation; Formal analysis; Investigation; Project administration.

Competing interests: No competing interests declared.

**Keni Vidilaseris**

Molecular and Integrative Biosciences, Biological and Environmental Sciences, University of Helsinki, 00100 Helsinki, Finland.

Contribution: Methodology; Validation; Formal analysis; Investigation; Writing – original draft preparation; Writing – review & editing; Visualization.

Competing interests: No competing interests declared.

**Orquidea Ribeiro**

Molecular and Integrative Biosciences, Biological and Environmental Sciences, University of Helsinki, 00100 Helsinki, Finland.

Contribution: Investigation.

Competing interests: No competing interests declared.

**Jianing Liu**

Molecular and Integrative Biosciences, Biological and Environmental Sciences, University of Helsinki, 00100 Helsinki, Finland.

Contribution: Validation; Investigation; Visualization. Competing interests: No competing interests declared. James Hillier

Astbury Centre for Structural and Molecular Biology, University of Leeds, LS2 9JT, Leeds, UK. Contribution: Investigation.

Competing interests: No competing interests declared.

**Anssi Malinen**

Department of Life Technologies, University of Turku, FIN-20014 Turku, Finland

Contribution: Methodology; Validation; Formal analysis; Investigation; Resources; Writing – review & editing.

Competing interests: No competing interests declared.

**Bernadette Gehl**

Molecular and Integrative Biosciences, Biological and Environmental Sciences, University of Helsinki, 00100 Helsinki, Finland.

Contribution: Investigation.

Competing interests: No competing interests declared.

**Lars J.C. Jeuken**

Leiden Institute of Chemistry, University Leiden, PO Box 9502, 2300 RA Leiden, Netherlands.

Contribution: Methodology; Validation; Formal analysis; Resources; Writing – review & editing; Supervision.

Competing interests: No competing interests declared.

**Arwen R. Pearson**

Institute for Nanostructure and Solid State Physics, Hamburg Centre for Ultrafast Imaging, Universität Hamburg, 22761 Hamburg, Germany

Contribution: Conceptualization; Methodology; Formal analysis; Investigation; Resources; Writing – review & editing; Supervision; Project administration; Funding acquisition.

Competing interests: No competing interests declared.

**Adrian Goldman**

Astbury Centre for Structural and Molecular Biology, University of Leeds, LS2 9JT, Leeds, UK and Molecular and Integrative Biosciences, Biological and Environmental Sciences, University of Helsinki, 00100 Helsinki, Finland.

Contribution: Conceptualization; Methodology; Validation; Formal analysis; Resources; Writing – original draft preparation; Writing – review & editing; Supervision; Project administration; Funding acquisition.

Competing interests: No competing interests declared.

## Data availability

All data needed to evaluate the conclusions in the paper are presented here. Additional data related to this paper may be requested from the authors. The atomic coordinates and structure factors of the *Pa*PPase:Mg_5_IDP complex (PDB ID: 8B37) and the grouped/combined time-resolved *Tm*PPase structures at 0-60-seconds (PDB ID: 8B21), 300-seconds (PDB ID: 8B22), 600-seconds (PDB ID: 8B23) and 3600-seconds (PDB ID: 8B24) have been deposited in the PDB, www.rcsb.org.

